# MultiMAP: Dimensionality Reduction and Integration of Multimodal Data

**DOI:** 10.1101/2021.02.16.431421

**Authors:** Mika Sarkin Jain, Krzysztof Polanski, Cecilia Dominguez Conde, Xi Chen, Jongeun Park, Lira Mamanova, Andrew Knights, Rachel A. Botting, Emily Stephenson, Muzlifah Haniffa, Austen Lamacraft, Mirjana Efremova, Sarah A. Teichmann

## Abstract

Multimodal data is rapidly growing in many fields of science and engineering, including single-cell biology. We introduce MultiMAP, an approach for dimensionality reduction and integration of multiple datasets. MultiMAP recovers a single manifold on which all of the data resides and then projects the data into a single low-dimensional space so as to preserve the structure of the manifold. It is based on a framework of Riemannian geometry and algebraic topology, and generalizes the popular UMAP algorithm^1^ to the multimodal setting. MultiMAP can be used for visualization of multimodal data, and as an integration approach that enables joint analyses. MultiMAP has several advantages over existing integration strategies for single-cell data, including that MultiMAP can integrate any number of datasets, leverages features that are not present in all datasets (i.e. datasets can be of different dimensionalities), is not restricted to a linear mapping, can control the influence of each dataset on the embedding, and is extremely scalable to large datasets. We apply MultiMAP to the integration of a variety of single-cell transcriptomics, chromatin accessibility, methylation, and spatial data, and show that it outperforms current approaches in preservation of high-dimensional structure, alignment of datasets, visual separation of clusters, transfer learning, and runtime. On a newly generated single-cell Assay for Transposase-Accessible Chromatin using sequencing (scATAC-seq) and single-cell RNA-seq (scRNA-seq) dataset of the human thymus, we use MultiMAP to integrate cells along a temporal trajectory. This enables the quantitative comparison of transcription factor expression and binding site accessibility over the course of T cell differentiation, revealing patterns of transcription factor kinetics.

## Introduction

Multimodal data is rapidly growing in many fields of science and engineering, including single-cell biology. Emerging single-cell technologies are providing high-resolution measurements of different features of cellular identity, including single-cell assays for gene expression, protein abundance^2,3^, chromatin accessibility^4^, DNA methylation^5^, and spatial resolution^6^. Large scale collaborations including the Human Cell Atlas international consortium^7,8^ are generating an exponentially increasing amount of data of many biological tissues, using a myriad of technologies. Each technology provides a unique view of cellular biology and has different strengths and weaknesses. Integrating these measurements in the study of a single biological system will open avenues for more comprehensive study of cellular identity, cell-cell interactions, developmental dynamics, and tissue structure^9^.

The integration of multi-omic data poses several challenges^10^. Different omics technologies measure distinct unmatched features with different underlying distributions and properties and hence produce data of different dimensionality. This makes it difficult to place data from different omics in the same feature space. Additionally, omics technologies can also have different noise and batch characteristics which are challenging to identify and correct. Further, as multi-omic data grows along two axes, the number of cells per omic and the number of omics per study, integration strategies need to be extremely scalable.

Most data integration methods project multiple measurements of information into a common low-dimensional representation to assemble multiple modalities into an integrated embedding space. Recently published methods employ different algorithms to project multiple datasets into an embedding space, including canonical correlation analysis (CCA)^11^, nonnegative matrix factorization (NMF)^12^ or variational autoencoders^13^. In the field of genomics, single-cell transcriptomics, as a well-established method, often serves as a common reference, facilitating the transfer of cell type annotation and data across multiple technologies and modalities. While these methods can be tremendously powerful, they require correspondence between the features profiled across omics technologies. Another limitation of many existing methods is they are challenged by scaling to large datasets.

Here we introduce a method that overcomes all these limitations: MultiMAP, an approach for the dimensionality reduction and integration of multiple datasets. MultiMAP integrates data by constructing a non-linear manifold on which diverse high-dimensional data reside and then projecting the manifold and data into a shared low-dimensional space. In contrast to other integration strategies for single-cell data, MultiMAP can integrate any number of datasets, is not restricted to a linear mapping, leverages features that are not present in all datasets (*i*.*e*. datasets can be of different dimensionalities), can control the influence of each dataset on the embedding, and is effortlessly scalable to large datasets. The ability of MultiMAP to integrate datasets of different dimensionalities allows the strategy to leverage information that is not considered by methods that operate in a shared feature space. (e.g. MultiMAP can integrate the 20,000-feature gene space of scRNAseq data together with a 100,000-feature peak space of scATACseq data).

We apply MultiMAP to challenging synthetic multimodal data, and demonstrate its ability to integrate a wide range of single-cell omics datasets. Finally, we apply the approach to the study of T cell development with new scATACseq data from fetal thymi. We show that MultiMAP can co-embed datasets across different technologies and modalities, while at the same time preserving the structure of the data, even with extensive biological and technical differences. The resulting embedding and shared neighborhood graph (MultiGraph) can be used for simultaneous visualisation and integrative analysis of multiple datasets. With respect to single cell genomics data, this allows for standard analysis on the integrated data, such as cluster label transfer, joint clustering, and trajectory analysis.

## Results

### The MultiMAP Framework

We introduce MultiMAP, an approach for integration and dimensionality reduction of multimodal data based on a framework of Riemannian geometry and algebraic topology. MultiMAP takes as input any number of datasets of potentially differing dimensions. MultiMAP recovers geodesic distances on a single latent manifold on which all of the data is uniformly distributed. The distances are calculated between data points of the same dataset by normalizing distances with respect to a neighborhood distance specific to the dataset, and between data points of different datasets by normalizing distances between the data in a shared feature space with respect to a neighborhood parameter specific to the shared feature space. These distances are then used to construct a neighborhood graph (MultiGraph) on the manifold. Finally, the data and manifold space are projected into a low-dimensional embedding space by minimizing the cross entropy of the graph in the embedding space with respect to the graph in the manifold space. MultiMAP allows the user to modify the weight of each dataset in the cross entropy loss, allowing the user to modulate the contribution of each dataset to the layout. Integrated analysis can be performed on the embedding or the graph, and the embedding also provides an integrated visualization. The mathematical formulation of MultiMAP is elaborated in Supplementary Methods.

In order to study MultiMAP in a controlled setting, we first applied it to two synthetic examples of multimodal data (Methods). The first synthetic data consists of points sampled randomly from the canonical 3D “Swiss Roll” surface and the 2D rectangle (Figure 2a). The dataset is considered multimodal data, because samples are drawn from different feature spaces but describe the same rectangular manifold. In addition, we are given the position along the manifold of 1% of the data. This synthetic setting illustrates that MultiMAP can integrate data in a nonlinear fashion and operate on datasets of different dimensionality, because data points along a similar position on the manifold are near each other in the embedding (Figure 2b). The MultiMAP embedding properly unrolls the Swiss Roll dataset, indicating that the projection is nonlinear. The embedding also appears to preserve aspects of both datasets; the data is curved and at the same time unrolled.

To determine if MultiMAP can effectively leverage features unique to certain datasets, we used the MNIST database^14^, where handwritten images were split horizontally with thin overlap (Figure 2c; see Methods for details). The two datasets can be considered multimodal because they have different feature spaces but describe the same set of digit images. The thin overlapping region of the two halves is not enough information to create a good embedding of the data (Figure 2c). Many distinct digits are similar in this thin central sliver, and hence they cluster together in the feature space of this sliver. Indeed, in a UMAP projection of the data in the shared feature space of this overlap, the clusters of different digits are not as well separated as in the UMAP projections of each half (Figure 2c).

A multimodal integration strategy that effectively leverages all features would use the features unique to each half to separate different digits, and the shared space to bring the same digits from each dataset close together (Figure 2d). We show that with MultiMAP the different modalities are well mixed in the embedding space and the digits cluster separately, despite mostly different feature spaces and noise being added to only the second dataset. This indicates that MultiMAP is leveraging the features unique to each dataset and is also robust to datasets with different noise.

Moreover, MultiMAP has weight parameters ω^v^ which control the contribution of each dataset **X**^v^ to the final embedding, allowing the user to modulate which dataset has a greater influence on the MultiMAP embedding. When a dataset’s weight is larger, its structure has a larger contribution to the MultiMAP embedding. Our results show that when integrating the MNIST data, for different choices of ω^v^, the datasets remain well integrated in the embedding space (Extended Data Figure 1a,b).

**Figure 1.**
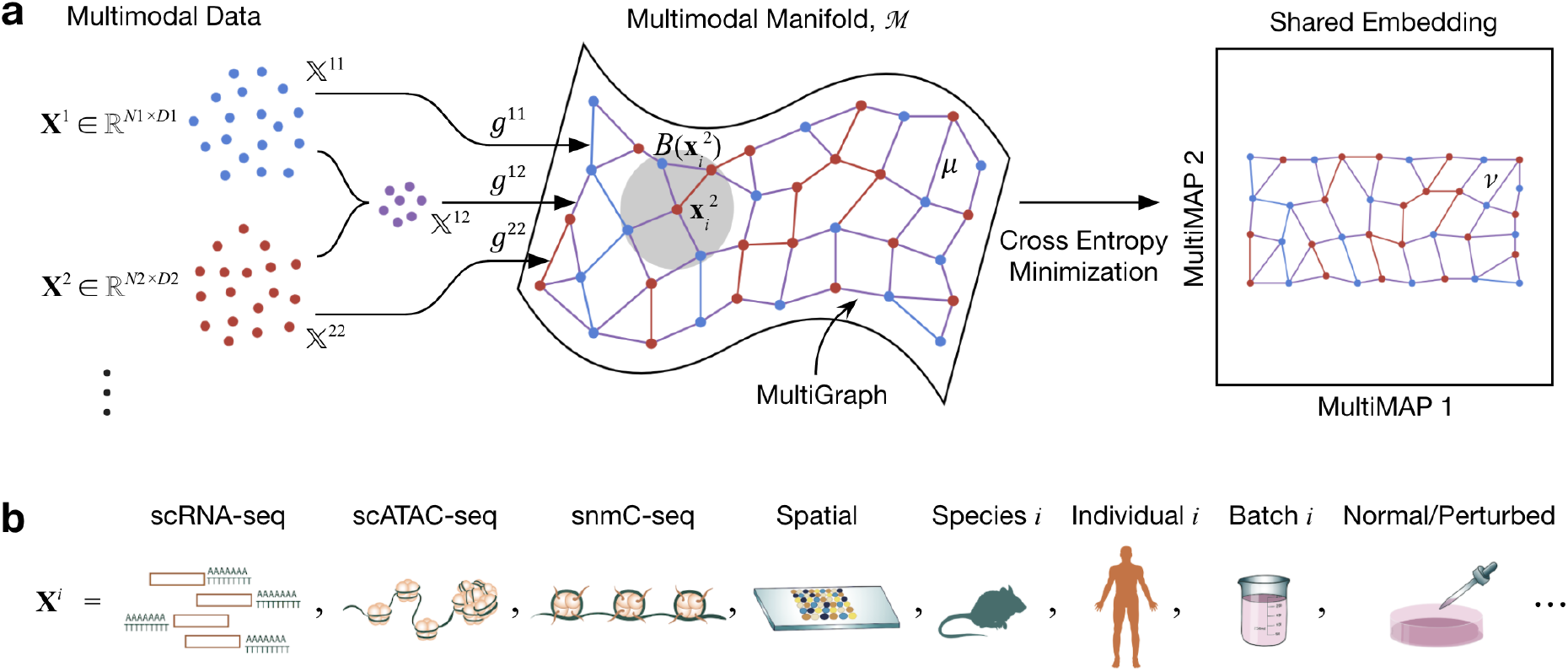
Schematic of MultiMAP. **a**. MultiMAP takes any number of datasets, including those of differing dimensions, recovers geodesic distances on a single latent manifold on which all data lie, constructs a neighborhood graph (MultiGraph) on the manifold, and then projects the data into a single low-dimensional embedding. Integrated analysis and visualisation can be performed on the embedding or graph. Variables are discussed in Methods. **X**^i^ is dataset *i*, **x**_j_^i^ is a point in **X**^i^, M is the shared manifold, *B*(**x**_i_^2^) is a ball on M centered at **x**_i_^2^, X^ij^ is the ambient space of M in the coordinate space with data containing points from datasets *i* and *j, g*^ij^ is the metric of M in the space X^ij^, *μ* is the membership function of the fuzzy simplicial set on the manifold, *ν* is the membership function of the fuzzy simplicial set in the low-dimensional space. **b**. In the field of cell atlas technologies, encompassing single cell genomics and spatial technologies, MultiMAP can be applied to integrate across different omics modalities, species, individuals, batches, and normal/perturbed states.

**Figure 2.**
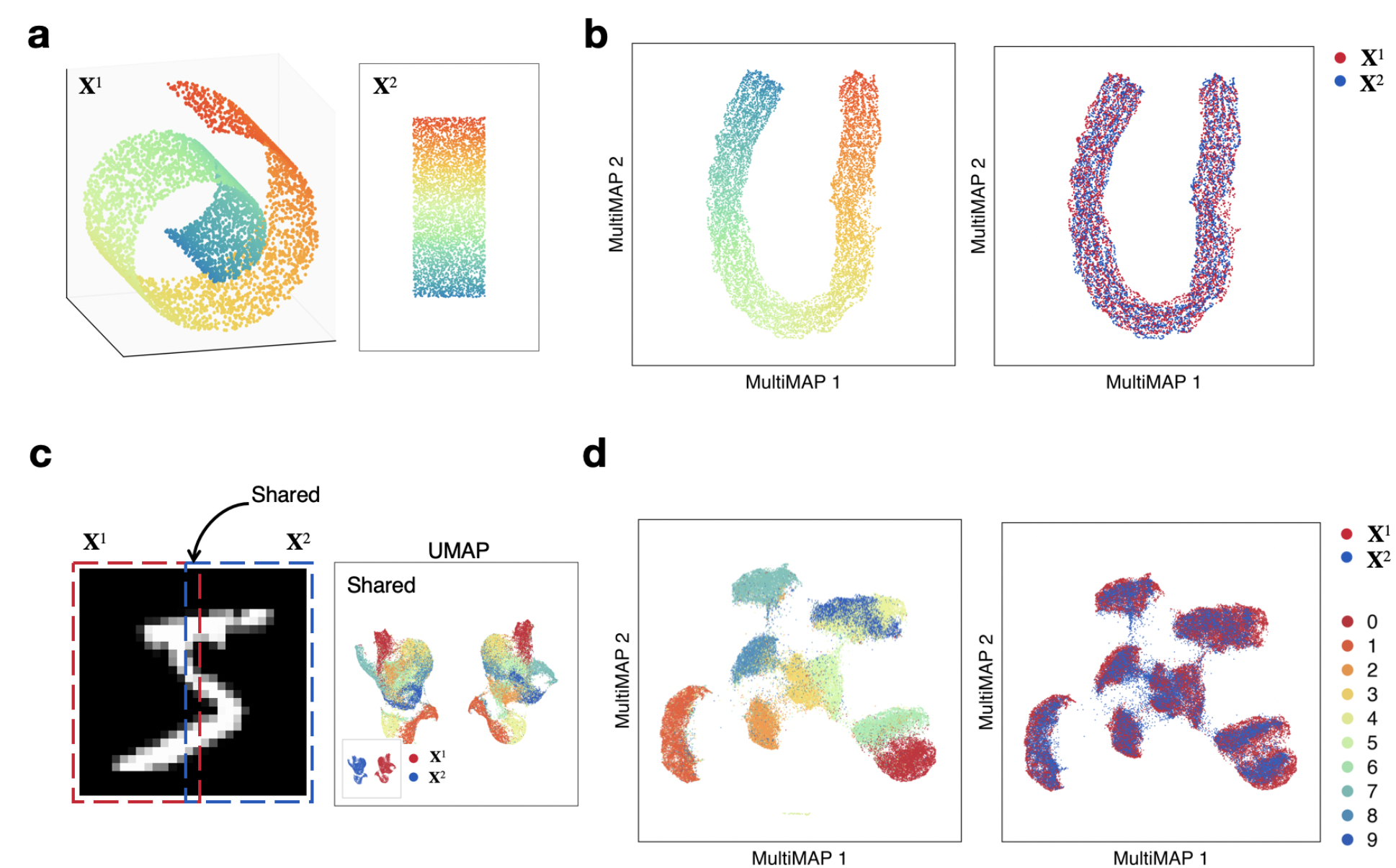
MultiMAP applied to synthetic data. **a**. Data sampled from the 3D Swiss Roll (**X**^1^) and a 2D rectangle (**X**^2^). **b**. Shared embedding of both datasets produced by MultiMAP. Color indicates position along the manifold (a,b). **c**. Left (**X**^1^) and right (**X**^2^) halves of MNIST handwritten digit images with a 2 pixel wide shared region. Gaussian noise is added to the left half. UMAP projections of each half and the shared region. **d**. Shared embedding of both MNIST halves (including Gaussian noise introduced for the left half) produced by MultiMAP. Each color is a different handwritten digit (0-9 as shown in the key). This illustrates that MultiMAP leverages both shared and unshared features to integrate multimodal datasets.

Finally, to illustrate that our assumption of a shared manifold is robust to variable levels of overlap across datasets, we used MultiMAP to integrate datasets with varying numbers of shared clusters in the MNIST data (Extended Data Figure 2). Our results show that MultiMAP is able to effectively integrate datasets that have only 1 out of 10 clusters shared between them. The transfer accuracy, silhouette score, and structure score of the MultiMAP integration remained largely constant as the number of overlapping clusters is varied, demonstrating that MultiMAP is highly robust to differences in populations between datasets.

## MultiMAP integration of single-cell transcriptomics and chromatin accessibility

Having shown that MultiMAP succeeds in integrating synthetic data, we apply the technique to real biological data. Epigenomic regulation underlies gene expression and cellular identity. Hence, integration of single-cell transcriptomics and epigenomics data provides an opportunity to investigate how epigenomic alterations regulate gene expression to determine and maintain cell identity. In addition, effective integration with transcriptomics data can improve the sensitivity and interpretability of the sparse scATAC-seq data.

To assess MultiMAP’s ability to integrate transcriptomic and epigenomic data, we applied the approach to integrate our previously generated high-coverage scATAC-seq data of mouse splenocytes^15^ and generated corresponding single-cell transcriptomic profiles of the same tissue. The high coverage of the plate-based scATAC-seq data as well as the published cluster annotations of the subpopulations served as a good ground truth example to validate our method. The analysis of the transcriptomics data revealed similar subpopulations to the published scATAC-seq dataset, in addition to two RNA-specific clusters: a subpopulation of B cells with higher expression of Interferon-Induced (Ifit) genes and a subpopulation of proliferating cells (Extended Data Figure 3a,b).

**Figure 3.**
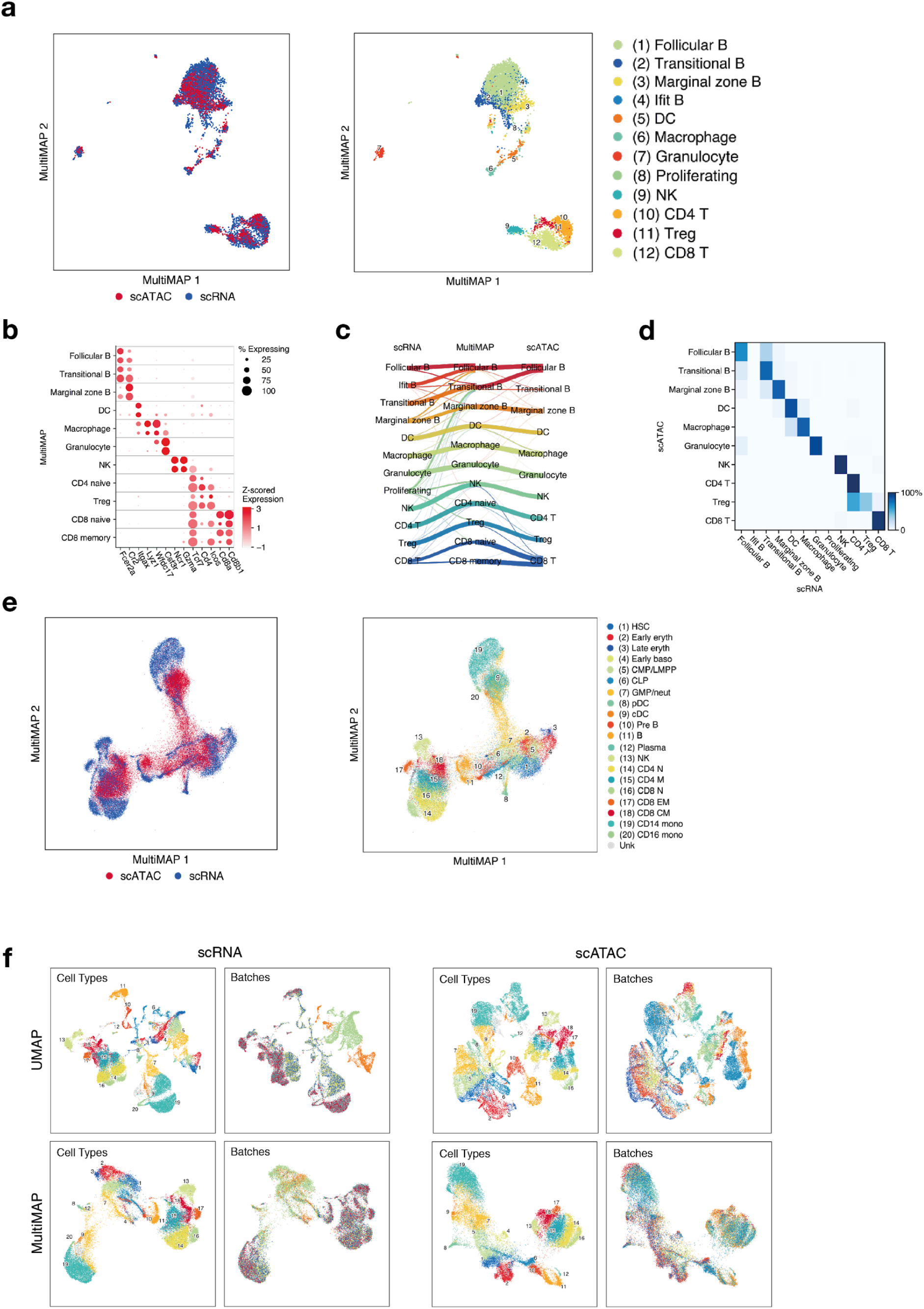
MultiMAP integration of single-cell transcriptomics and chromatin accessibility. **a**. MultiMAP visualization of the integration of published scATAC-seq^15^ and newly generated scRNA-seq data of the mouse spleen (n=1), colored by omic technology (left hand panel) and independent cell type annotations of each omic technology (right hand panel). **b**. Dot plot showing the z-score of the mean log-normalised gene expression and gene activity scores of known markers of each identified joint cluster. The top dot of each row shows the cells from the scRNA-seq data, and the bottom dot represents the cells from the scATAC-seq data. **c**. Riverplot showing correspondence between the joint clusters and the independent annotations of the scATACseq and scRNAseq data. **d**. Confusion matrix of label transfer from the scRNAseq to the scATACseq. **e**. MultiMAP visualization of the integration of single-cell transcriptomics and chromatin accessibility of human bone marrow and peripheral blood mononuclear cells^16^ colored by omic technology (left hand panel) and by the published cell type annotation (right hand panel). **f**. UMAP (panels in top row) and MultiMAP (panels in bottom row) visualization of the scRNA-seq and scATAC-seq data colored by cluster annotation and batch, showing the effective batch correction of both modalities using MultiMAP.

MultiMAP effectively integrated the two datasets, using both gene activity scores and the cell type-specific epigenetic information outside of gene bodies. The different modalities are well mixed in the embedding space and cells annotated as the same type are close together, regardless of the modality for different choices of ω^v^ (Figure 3a, Extended Data Figure 1c,d). Next, we jointly clustered cells from both datasets using the MultiGraph. This produced clusters with markers corresponding to known cell types^15^ (Extended Data Figure 3c). The annotations produced by this joint clustering were generally consistent with independent annotations of each dataset (Figure 3c). Two of the clusters determined to be proliferating cells and B cells with upregulated Ifit genes were found only in the scRNA-seq data, as expected (Figure 3a, Extended Data Figure 3b). In addition, the integration produced by MultiMAP is robust to different choices of the weight parameters (Extended Data Figure 1c).

Further, we used the MultiGraph to directly predict the cell types of the scATAC-seq given the cell types of the scRNA-seq. Figure 3d shows the confusion matrix of the predictions, illustrating that cells were generally annotated correctly. This illustrates the ability of MultiMAP to leverage annotation efforts of one omic technology to inform those of another. Interestingly, a small subset of cells from scRNA-seq previously annotated as T cells is now clearly separated on the MultiMAP plot, and clusters close to the B cells (Figure 3a, Extended Data Figure 3). Doublet detection confirmed that this cluster is composed of doublet T/B cells. These doublets are spread throughout the UMAP plot of the scRNA-seq data, but are clearly distinct on the MultiMAP plot (Extended Data Figure 3). This illustrates the power of MultiMAP both as a visualization tool, and to reveal new populations of cells.

Next, we applied MultiMAP to integration of multiple batches from each data modality, to assess the ability to account for batch effects. For this purpose, we used recently published scRNA-seq and scATAC-seq data of human bone marrow and peripheral blood mononuclear cells^16^. This dataset consists of 16 experimental samples, representing different experimental batches. Another challenge is that cells are not in discrete clusters but rather on a continuum. MultiMAP is able to simultaneously correct batch effects and modality differences, integrating all 16 datasets into a consistent embedding (Figure 3e). The different modalities are well mixed in the embedding and cells of the same type are close together, regardless of modality or batch. The cell type annotations of all of the data were taken from the original publication^16^, so they provide a good ground truth and independent validation of MultiMAP. Additionally, MultiMAP is able to correct batch effects present in different omics technologies. Applying MultiMAP to just the scRNA-seq data produces embedding that properly integrates cells of the same type regardless of batch, and the same is true when MultiMAP is applied to only the scATAC-seq data (Figure 3f). It is also evident in this figure that clusters with cell types unique to a batch remain unmixed in the embedding. This indicates that MultiMAP is not forcing incompatible data to integrate and demonstrates that MultiMAP can integrate datasets even if they have extensive technical differences.

### MultiMAP integration of multiple modalities of mouse brain cells

Recent advances in spatial sequencing technology enable the simultaneous measurement of gene expression and spatial locations of single-cells, facilitating the study of tissue structure^6^. While these technologies provide spatial information, they often measure only a small fraction of the genes measured by scRNA-seq. Integration of spatial measurements and scRNA-seq has the potential to provide spatial context to scRNA-seq data as well as to reveal finer grained biological differences in the spatial data by leveraging the greater number of cells and genes present in scRNA-seq data.

We applied MultiMAP to the integration of a Drop-seq scRNA-seq data of the mouse frontal cortex^17^ and STARmap *in situ* gene expression dataset^18^. Despite the differences between the two dataset in the number of measured genes (only 1020 in STARmap) and the number of cells (71640 in Drop-seq versus 2137 in STARmap), our integrated analysis shows that MultiMAP successfully integrates the datasets. Clustering the integrated data using the MultiGraph produced clusters with markers corresponding to known cell types (Figure 4a,b). One of the clusters, the claustrum, was found only in the scRNA-seq data, as expected. Integration with MultiMAP also resulted in improved cluster annotation for both datasets. The excitatory L4 neurons were previously only present in the STARMap data, as the motor cortex and prefrontal cortex that are part of the frontal cortex are considered to lack a layer 4 in mice^19^. However, after the integration we also identified L4 cells in the scRNA-seq data previously annotated as L5 neurons (Figure 4a,c, Extended Data Figure 4). A similar population of pyramidal cells located between layers 3 and 5 were recently identified both with anatomical and single-cell studies ^20,21^. This was confirmed by expression of marker genes associated with L4, including Cux2 and Rorb (Extended Data Figure 4). This illustrates the power of MultiMAP to reveal new cell types.

**Figure 4.**
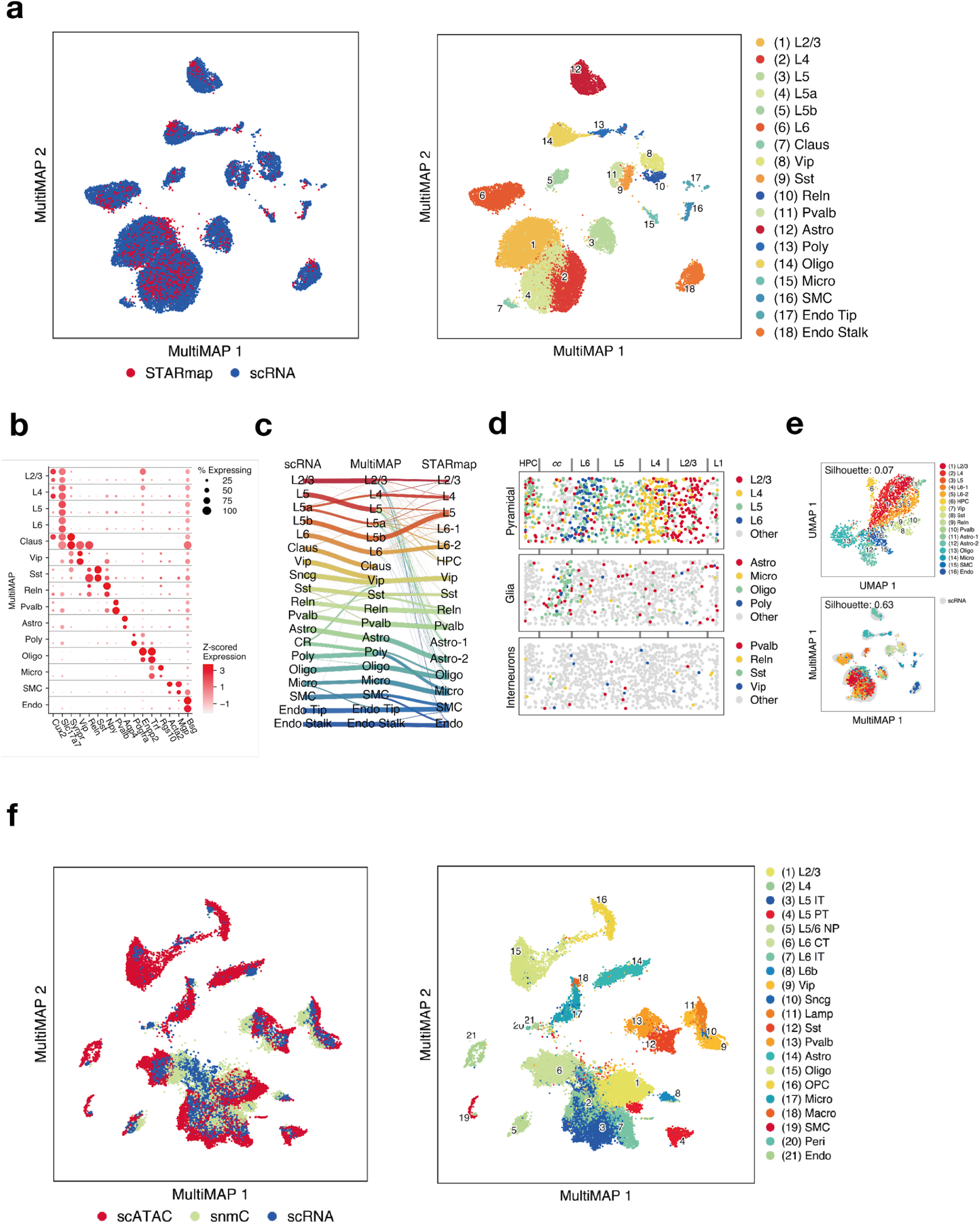
MultiMAP integration of multiple modalities of mouse brain cell data. **a**. MultiMAP visualization of scRNA-seq^17^ (n=2) and spatial STARmap^18^ (n=2) data of the mouse brain, colored by omic technology and joint clusters identified with the MultiGraph. **b**. Dot plot showing mean log-normalised gene expression of known markers of each identified joint cluster. The top dot in each row represents cells from the scRNA-seq data, and the bottom dot represents cells from the scATAC-seq data. **c**. Riverplot showing correspondence between the joint clusters, and the independent annotations of the scATACseq and scRNAseq data. **d**. Spatial locations of the STARmap cell, colored by the joint clusters. **e**. UMAP and MultiMAP visualizations of the STARmap dataset. The silhouette score as employed here quantifies the separation of clusters, and the higher value for MultiMAP shows the better cluster separation as compared to UMAP. **f**. MultiMAP visualization of the integration of single-cell transcriptomics, chromatin accessibility, and DNA methylation of the mouse primary cortex, colored by omic technology and the published cell type annotation^20^.

MultiMAP also improves visualization of the STARmap data. Before integration with MultiMAP, many of the cell types of the spatial data did not cluster separately and were visually hard to distinguish. In comparison, the MultiMAP embedding of the STARmap data exhibits tighter cell type clusters and increased separation between cell types (Figure 4e). This improvement was measured by the average Silhouette score in the embedding space, which is significantly larger for MultiMAP (Figure 4e).

Integration with MultiMAP also enabled us to spatially locate all the joint cell types in the STARmap data, allowing study of the spatial structure of the tissue (Figure 4d). The pyramidal neurons localize to layers 2-6 and oligodendrocytes localize to the layer below the cortex, whereas the interneurons do not appear to exhibit spatial organization. These observations are all consistent with the known spatial architecture of the mouse visual cortex^18^.

To investigate the performance of MultiMAP on the integration of more than two modalities, we applied the approach to integrate recently published multi-omics datasets of the mouse primary motor cortex^20^ consisting of 9 separate datasets, including 7 single-cell or single-nucleus transcriptomics datasets, one single-nucleus chromatin accessibility, and one single-nucleus DNA methylation (snmC-seq). MultiMAP successfully co-embedded more than 600,000 single-cell or -nucleus samples assayed by six molecular modalities and identified the previously published cell subpopulations. The MultiMAP embedding displays good mixing of clusters from different modalities when the clusters correspond to the same cell type. Cell type annotations were taken from the original publication of the data, so they provide a good ground truth and an independent validation of MultiMAP. We further see that cell types that exist in one modality, but not in the others, are not falsely aligned in the embedding space. This indicates that MultiMAP is not forcing incompatible data to integrate.

Finally, using the integration of scRNA-seq with the STARmap data, as well as the integration of the multi-omics spleen data, we assessed the impact of using only shared vs. all features. We find that using all features greatly improves the integration and results in embeddings that are visually and quantitatively superior, according to four performance metrics (Extended Data Figure 5). This illustrates that non-shared features can be extremely helpful, and demonstrates an advantage of MultiMAP over other methods which do not consider non-shared features.

**Figure 5.**
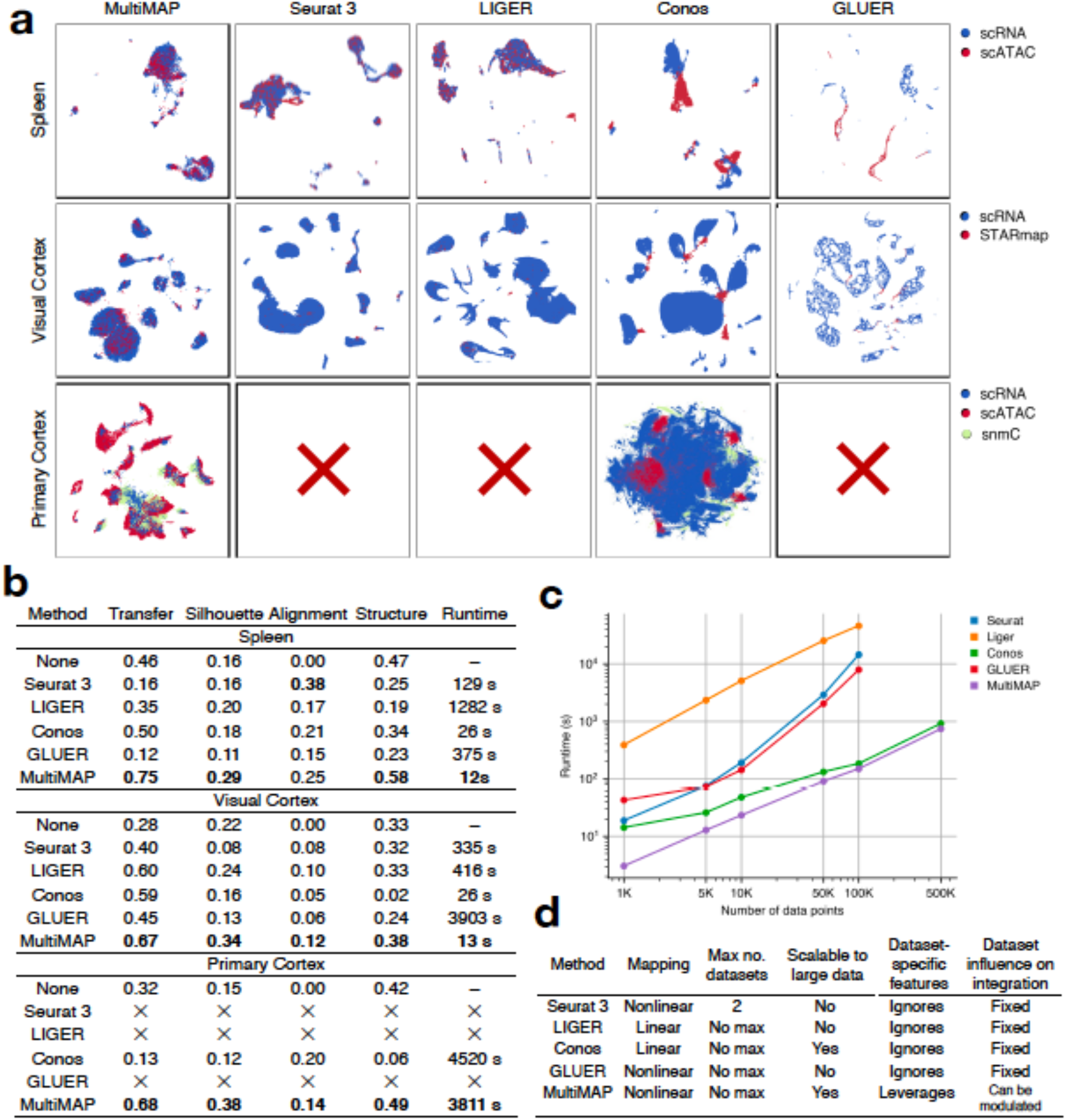
Benchmarking MultiMAP against existing approaches. **a**. Embeddings returned by multi-omic integration methods on different datasets. “X” indicates that the method terminated due to an out-of-memory error (218 GB RAM). **b**. Comparison of each method in terms of transfer learning accuracy (“Transfer”), separation of cell type clusters as quantified by Silhouette coefficient (“Silhouette”), mixing of different datasets as measured by fraction of nearest neighbours that belong to a different dataset (“Alignment”), preservation of high-dimensional structure as measured by the Pearson correlation between distances in the high-and low-dimensional spaces (“Structure”), and runtime. **c**. Wall-clock time of multi-omic integration methods on different sized datasets. Seurat 3 and LIGER produced out-of-memory errors when run on 500,000 data points (218 GB RAM). To produce these datasets we subsampled the mouse primary cortex scRNA-seq and scATAC-seq data^20^ using geometric sketching^33^. The datasets were subsampled so that there are equal number of cells in the scRNA-seq and scATAC-seq data until 100,000 cells. Since the scATAC-seq data had 81,196 cells in total, for the 500,000 cells comparison, we used an scRNA-seq of 418,804 cells. **d**. Comparison of capabilities and properties of each method. “Mapping” refers to the nature of the mapping employed by the method; “Max no. datasets” refers to the upper limit in terms of numbers of datasets accepted by the method; “Scalable to large data” refers to allowing a total of over 500,000 cells; “Data-set specific features” is whether the integration method allows information that is not shared across datasets; and “Dataset influence on integration” is whether the user can modulate the weighting of a given dataset relative to the others during the integration.

### Benchmarking

We assessed and benchmarked the performance of MultiMAP against several popular approaches for integrating single-cell multi-omics, including Seurat 3^11^, LIGER^12^, Conos^22^ and GLUER^23^.

These integration approaches differ in key regards, summarized in Figure 5d. We used a diversity of performance metrics to comprehensively compare MultiMAP with other approaches, including transfer accuracy, silhouette score, alignment, preservation of the structure, and runtime. With these metrics, we quantified the separation of the joint clusters, how well mixed the datasets were after integration and how well they preserved the structure in the original datasets to investigate whether the methods integrate populations across datasets without blending distinct populations together.

To this end, we generated single-nucleus data from human Peripheral Blood Mononuclear Cells (PBMCs) using the Multiome ATAC + RNA kit. We obtained a PBMC atlas of 6,344 nuclei of high-quality ATAC + RNA profiles. We analysed and annotated the RNA and ATAC data separately, revealing all the known major PBMC types: CD14 and CD16 monocytes, cDCs and pDCs, naive and effector CD4 and CD8 T cells, Tregs, MAIT and gamma-delta T cells, NK and ILCs, naive and memory B cells and plasmablasts (Extended Data Figure 6a). Most cell types were well separated in both modalities with the exception of the NK and ILC clusters and the gamma-delta and the CD8 effector T cells that blended together in the ATAC data.

**Figure 6.**
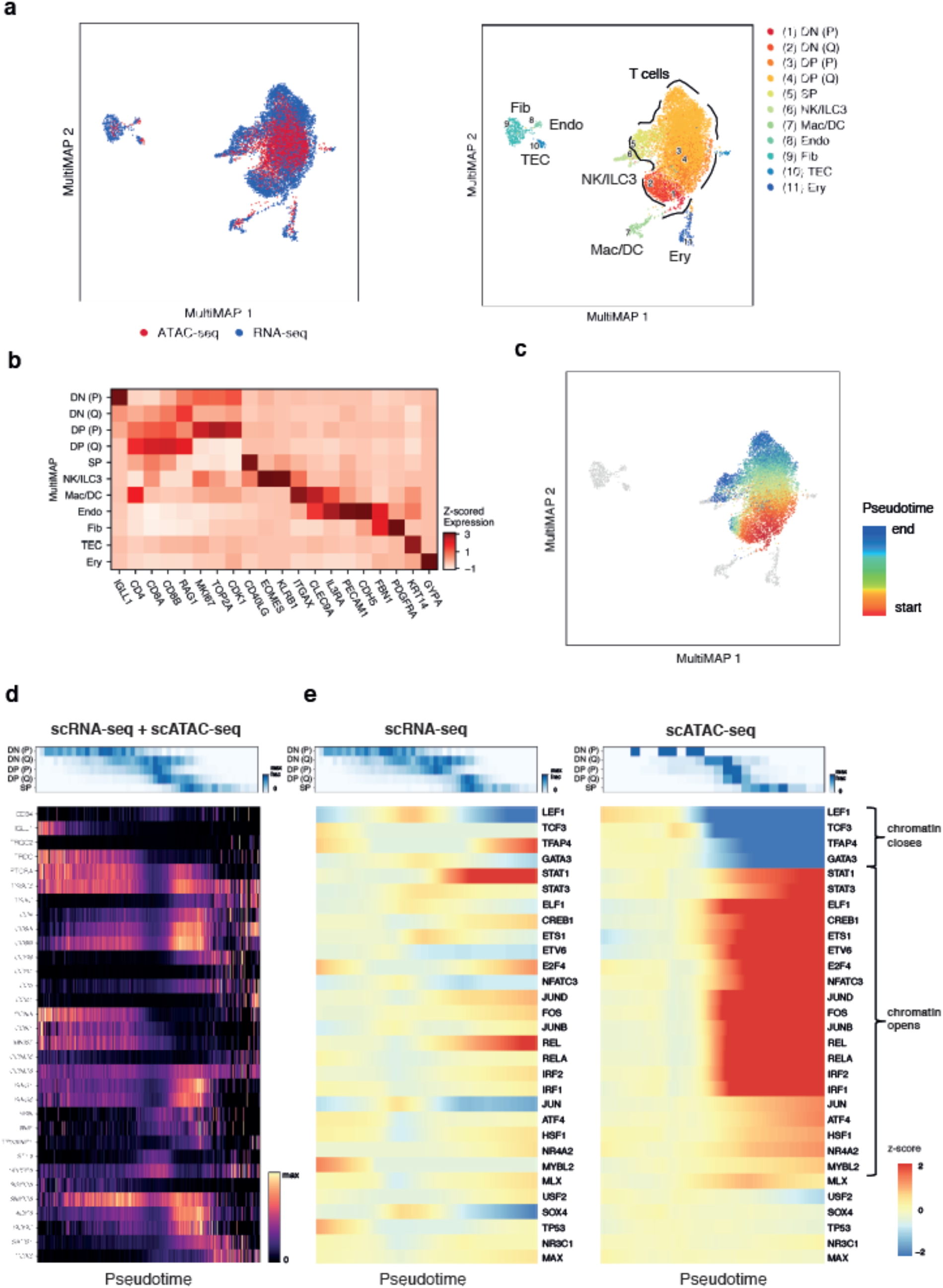
Integration of scRNAseq and scATACseq data of human fetal thymus reveals transcriptional regulatory principles of T cell development. **a**. MultiMAP visualization of scRNA-seq and scATAC-seq datasets of the human fetal thymus (n=1), colored by modality and joint clusters identified using the MultiGraph. **b**. Heatmap of gene expression and gene activity scores of key markers of the joint clusters identified using the MultiGraph. **c**. Inferred pseudotime using the MultiGraph recovers the T cell differentiation trajectory. Color indicates pseudotime from red (early, beginning) to blue (late, end). **d**. Heatmap of the gene expression and gene activity scores over pseudotime of genes known to be involved in T cell development. **e**. Smoothed heatmaps of the z-score of the gene expression and motif accessibility of the most variable transcription factors over pseudotime. The motif accessibilities of TFs that varied most in time show changes in accessibility at the transition between the late DN and early DP stage of differentiation. This includes TFs such as GATA3, ZEB1 for which the chromatin at the binding sites closes at that transition, and TFs for which the chromatin at the binding sites opens, such as E2F4, ETS1 and others.

We used the PBMCs as a gold standard dataset to benchmark MultiMAP against the four other methods. As shown in the co-embedding and the metrics, MultiMAP successfully integrated the cell types across modalities and outperformed other methods (Extended Data Figure 6b,c). The label transfer accuracy was particularly striking, with MultiMAP achieving a much higher score compared to other methods.

Furthermore, we also benchmarked MultiMAP using a variety of multi-omic data with published cell type annotations, including the transcriptomics and chromatin accessibility spleen data, scRNA-seq and STARmap of the visual cortex, and the multi-omics data of the primary cortex. For all datasets, MultiMAP achieves top or near top performance on all metrics (Figure 5a,b). The embeddings produced by MultiMAP prove superior for transferring cell type annotations between datasets, separating clusters of different cell populations, integrating datasets in a well-mixed manner, and capturing the high-dimensional structure of each dataset. Critically, MultiMAP is faster than all other benchmarked methods, and significantly faster than LIGER and Seurat 3 (Figure 5c). Seurat 3 and LIGER were not able to scale to the primary cortex data of 600k, producing out-of-memory errors despite access to 218 GB of RAM.

Finally, to assess the batch correction performance of MultiMAP, we also applied it on three scRNA-seq studies of the human pancreas^24–26^ that were recently used for comparison of eight batch correction methods^27^. Even though the main purpose of MultiMAP is the integration of several different omic technologies, MultiMAP outperformed all other well established batch correction methods in the field, demonstrating that MultiMAP can correct batches and integrate multiple omics data simultaneously (Extended Data Figure 7).

### MultiMAP reveals patterns of T cell maturation along a multi-omic trajectory

Single-cell transcriptomics has enabled reconstruction of developmental trajectories and the study of dynamic processes such as differentiation and reprogramming. Bulk RNA-seq and ATAC-seq data has further revealed regulatory events driving these processes^28^. However, joint analysis of single-cell expression and chromatin accessibility profiles along a time course trajectory would allow the study of dynamic chromatin regulation alongside gene expression and elucidate epigenomic drivers of transcriptional change^29,30^.

In order to investigate the potential of integrating multi-omic data along a common differentiation trajectory, we focused on T cell development in the thymus. The thymus is an organ essential for the maturation and selection of T cells. Precursor cells migrate from the fetal liver and bone marrow to the thymus where they develop into different types of mature T cells^31^. We recently provided a comprehensive single-cell transcriptomics atlas of the human thymus during development, childhood, and adult life, and computationally predicted the trajectory of T cell development from early progenitors to mature T cells^31^. To expand on this and further investigate the gene regulatory mechanisms driving T cell development, we generated single-cell transcriptomics and chromatin accessibility data from a human fetal thymus sample at 10 weeks of gestation.

Clustering of the scRNA-seq data revealed cell types identified in our recently published transcriptomic cell atlas of the thymus^31^, including several clusters of T cells across different stages of development, fibroblasts, endothelial cells, erythrocytes, thymic epithelial cells (TECs), NK and ILC3 cells, and macrophages and dendritic cells (Extended Data Figure 8). The sparse scATAC-seq and the continuous nature of cell types along the maturation trajectory made it difficult to cluster the ATAC cells into different T cell types (Extended Data Figure 8). However, the integration with MultiMAP and the joint clusters obtained using the MultiGraph corresponded to the published thymus cell types^31^ (Figure 6 a,b), allowing us to correctly annotate the cell types of the scATAC-seq data.

We then selected the T cell populations identified from the joint clustering and performed diffusion map pseudotime analysis using the alignment MultiMAP graph. The reconstructed development trajectory showed a continuous differentiation with the same trend as the published study, starting from early double negative (DN) CD4-CD8-, gradually progressing to double positive (DP) CD4+CD8+ T cells, and then differentiating into single positive (SP) mature CD8+ or CD4+ T cells. Hallmark genes of T cell differentiation varied along the inferred pseudotime in a manner consistent with^31^ (Figure 6d), serving as validation of the trajectory inference and the integration produced by MultiMAP.

To identify transcription factors (TFs) that potentially regulate T cell development, we studied changes in TF expression and TF binding site accessibility along the differentiation trajectory. The top variable TFs/TF binding sites along the trajectory included many TFs that have been previously shown to be involved in T cell differentiation, including GATA3, SPI1, MEF2C, ERG, TCF3, TCF4, TFAP4, MYBL2, STAT1, NR4A2 and others^28,31,32^ (Figure 6e, Extended Data Figure 9). The TFs that most varied along the trajectory were found to show changes in motif accessibility at the transition between the late DN and early DP stage of differentiation as shown before^32^.

Moreover, our integrated trajectory allowed us to identify TFs where changes in motif accessibility and expression of the TF itself were closely coordinated, for example ZEB1, IRF1, REL, FOS and others, suggesting that these TFs actively regulate their target genes immediately and directly (Figure 6e). In contrast, for TFs such as ETS1, JUN etc., gene expression of the TF significantly precedes the accessibility of the corresponding TF binding sites, suggesting that additional regulatory mechanisms are potentially required for opening of the TF motifs.

## Discussion

Here we present a novel approach for dimensionality reduction and integration of multimodal data which takes into account the full data sets, even when they have different feature spaces. MultiMAP embeds all datasets into a shared space to preserve both the manifold structure of each dataset independently, as well as in shared feature spaces. This enables both visualization and streamlined downstream analyses. Crucially, our method can incorporate different types of features, such as gene expression and open chromatin peaks or intergenic methylation, and thus takes advantage of the full power of multi-omics data. Ignoring the features unique to one dataset (as in most existing methods), may omit important information, for instance distinguishing features of certain subpopulations of cells and yield an integrated embedding that does not distinctly cluster all subpopulations.

An additional advantage of MultiMAP is that the influence of each dataset on the shared embedding can be modulated. This is useful when integrating datasets of different qualities, or when aligning a query dataset to a reference dataset. Comparison with existing methods for integration shows that MultiMAP outperforms or has close to best performance in every aspect investigated. MultiMAP is a robust and effective method for dimensionality reduction and integration of multimodal data, and is extremely fast and scalable to massive datasets.

Using synthetic examples to illustrate the power of the method, we show that MultiMAP leverages the features unique to each dataset to effectively integrate and reduce the dimensionality of the data, and is also robust to data with noise. Throughout our applications of MultiMAP to diverse single-cell multi-omic data, we demonstrate that our method can facilitate integration across transcriptomic, epigenomic, and spatially resolved datasets, and derive biological insights jointly from multi-omic single-cell data. In addition, our method can align datasets across different technologies and modalities even with extensive biological and technical differences. Crucially, we show that MultiMAP is flexible enough to integrate datasets with different clusters and cell populations, illustrating that MultiMAP is applicable even when its central hypothesis is not strictly reflected by the data. The multimodal integration of three or more omics technologies opens many opportunities for the comprehensive study of tissues.

We note that our method is based on the hypothesis that multi-omics data are uniformly distributed on a latent manifold. A hypothesis of this sort, about the distribution of data in a latent space, is a central feature of many existing integration strategies. For example, CCA-based strategies (including Seurat and Conos) assume that the data reside in a maximally-correlated manner in a latent space which is a linear projection of the original data. MultiMAP, in contrast, does not make as strong an assumption because we do not restrict the latent manifold to a linear projection of the data. While this kind of hypothesis is often realistic for data generated from the same tissue, there may be cases where this is not strictly the case. In practice, we find that MultiMAP can successfully accommodate datasets that depart from this central hypothesis, *i*.*e* when clusters and cell populations are not shared across all datasets that are being integrated.

Perhaps the greatest potential lies in applying MultiMAP to datasets beyond those considered here. Integrative analysis with MultiMAP can be used to compare healthy and diseased states, and identify pathologic features, or to uncover cell-type specific responses to perturbations. Other examples include the integration of data across species to enable studying the evolution of cell states and identifying conserved cell types and regulatory programs. Along similar lines, the integration of *in vivo* with *in vitro* models such as organoids will reveal the quality or faithfulness of cells in a dish relative to their native counterparts. Finally, given the rapid development of joint multimodal single cell genomics methods (e.g. CITEseq for protein and RNA, joint snRNA-and ATACseq), it is relevant to point out that MultiMAP can be applied to multi-omic data acquired both from different cells as well as from the same cells.

In summary, given the broad appeal of dimensionality reduction methods (e.g. PCA, tSNE, UMAP), and the growth of multimodal data in many areas of science and engineering, we anticipate that MultiMAP will find wide and diverse use.

## AUTHOR CONTRIBUTIONS

M.S.J., M.E. and S.A.T. conceived the study. M.S.J conceived and developed MultiMAP. M.S.J, created the codebase with contributions from M.E., and K.P. C.D.C., J.-E.P., R.A.B, E.S, L.M., A.E. and X.C. generated the single cell data. M.E. analysed the data and interpreted the results with contributions from M.S.J. and S.A.T. M.S.J., M.E. and S.A.T. wrote the manuscript with contributions from A.L., X.C., and C.D.C. All authors read and accepted the manuscript.

## DATA AVAILABILITY

scRNA-seq and scATAC-seq data generated for this publication were being deposited in ArrayExpress: E-MTAB-9769 for scRNA-seq of mouse splenocytes, E-MTAB-9840 and E-MTAB-9828 for scRNA-seq and scATAC-seq of the thymus (username; password: rmachcde). The Multiome RNA+ATAC PBMC data is in the process of being deposited.

## CODE AVAILABILITY

MultiMAP is publicly available at github.com/Teichlab/MultiMAP. ACKNOWLEDGEMENTS We thank Jana Eliasova for the graphical illustrations. We are grateful to Emma Dann, Natsuhiko Kumasaka and Zhihan Xu for critical feedback on the manuscript. We thank Ruben Chazarra-Gil and Vladimir Yu Kiselev for the comparison of different batch correction methods on the pancreas dataset. M.S.J. was supported by a Gates Cambridge Scholarship. J.-E.P. was supported by EMBO Long-Term and Advanced Fellowships. M.E. is funded by Barts Charity. S.A.T. is funded by Wellcome (WT206194). The study was supported by Wellcome Human Cell Atlas Strategic Science Support (WT211276/Z/18/Z) and the Chan Zuckerberg Initiative (CZF2019-002445).

## Supplementary Figures

**Extended Data Figure 1.**
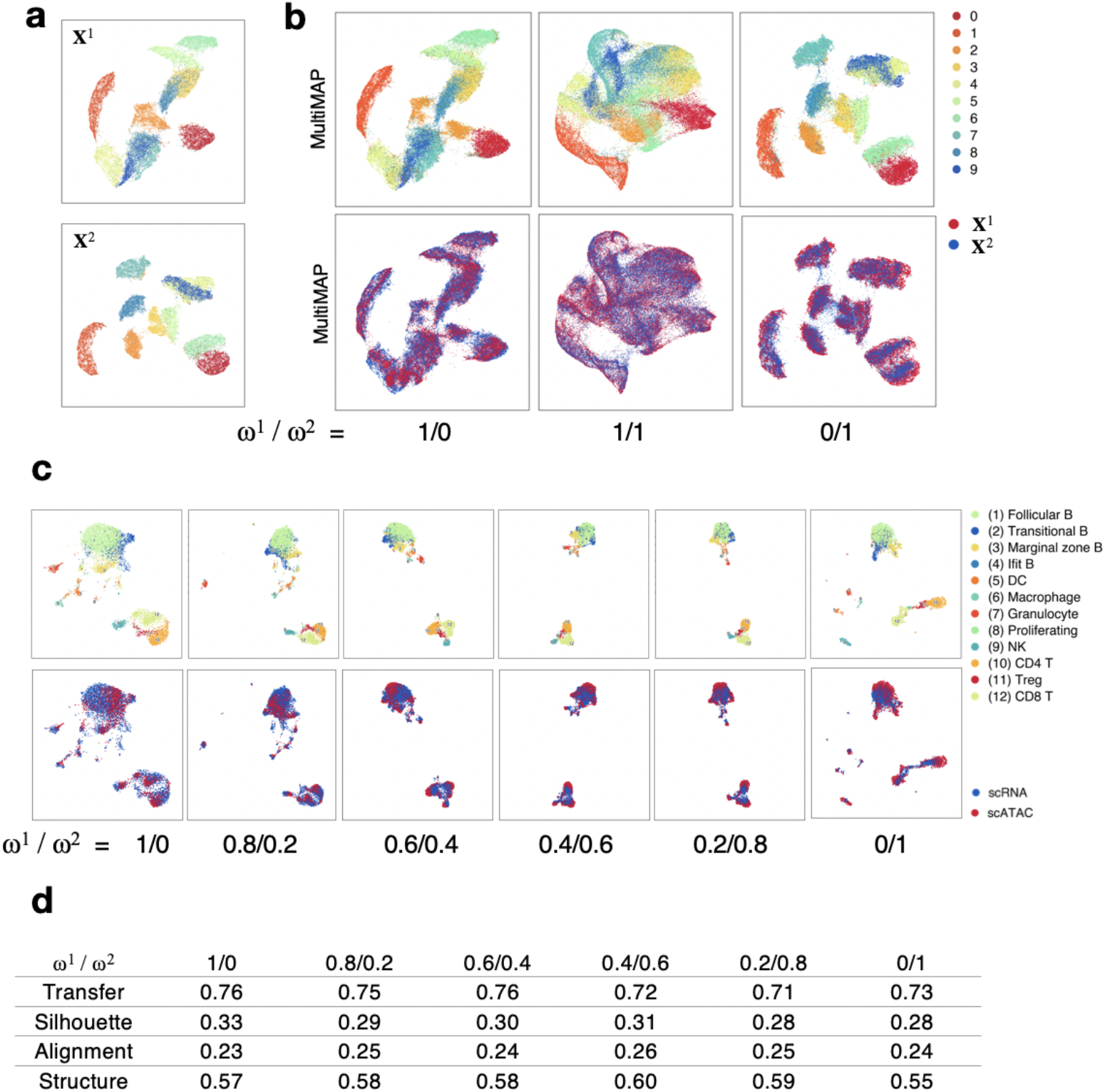
MultiMAP’s weight parameter. **a**. UMAP projections of the two halves of the MNIST handwritten digit images. **b**. MultiMAP embeddings as the weight parameters are varied. Each color is a different handwritten digit (0-9). When *ω*^1^ is larger than *ω*^2^, the embedding more closely resembles the projection of only **X**^1^; when *ω*^2^ is larger than *ω*^1^, the embedding more closely resembles the projection of only **X**^2^. For different choices of *ω*^v^, the datasets are well integrated in the embedding space. **c**. MultiMAP integration with varied weight parameters of published scATAC-seq^15^ and newly generated scRNA-seq data of the mouse spleen (n=1). **d**. Comparison of the MultiMAP integration of the spleen data as the weight parameter is varied --in terms of transfer learning accuracy (“Transfer”), separation of cell type clusters as quantified by Silhouette coefficient (“Silhouette”), and preservation of high-dimensional structure as measured by the Pearson correlation between distances in the high-and low-dimensional spaces (“Structure”)

**Extended Data Figure 2.**
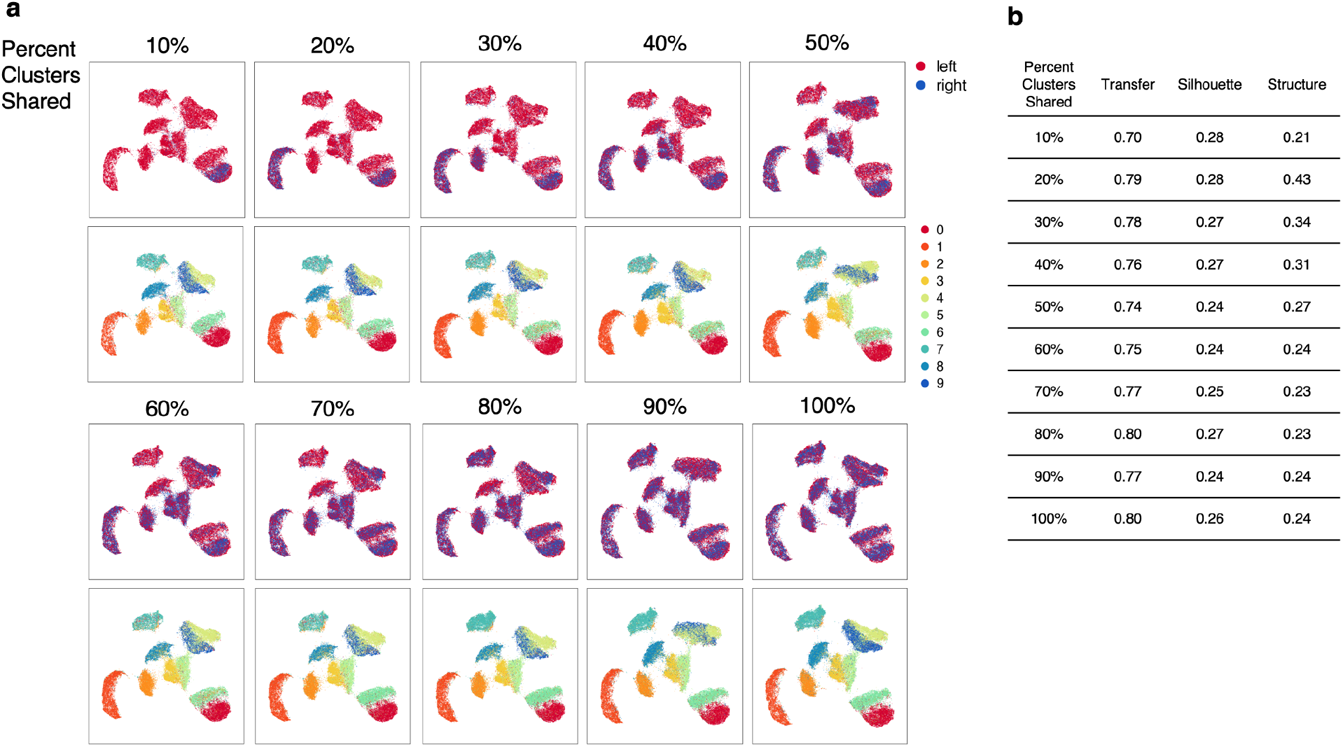
MultiMAP integration with non-shared clusters. **a**. MultiMAP integration of the left and right halves of MNIST handwritten digit images with a 2 pixel wide shared region. Gaussian noise is added to the left half. MultiMAP integration is performed with a varying number of digit clusters removed from the right dataset, so that the integration ranges from one shared cluster (10%) to all clusters shared (100%). **b**. Comparison of the MultiMAP integration of the modified MNIST dataset as the percent of clusters shared is varied --in terms of transfer learning accuracy (“Transfer”), separation of cell type clusters as quantified by Silhouette coefficient (“Silhouette”), and preservation of high-dimensional structure as measured by the Pearson correlation between distances in the high-and low-dimensional spaces (“Structure”).

**Extended Data Figure 3.**
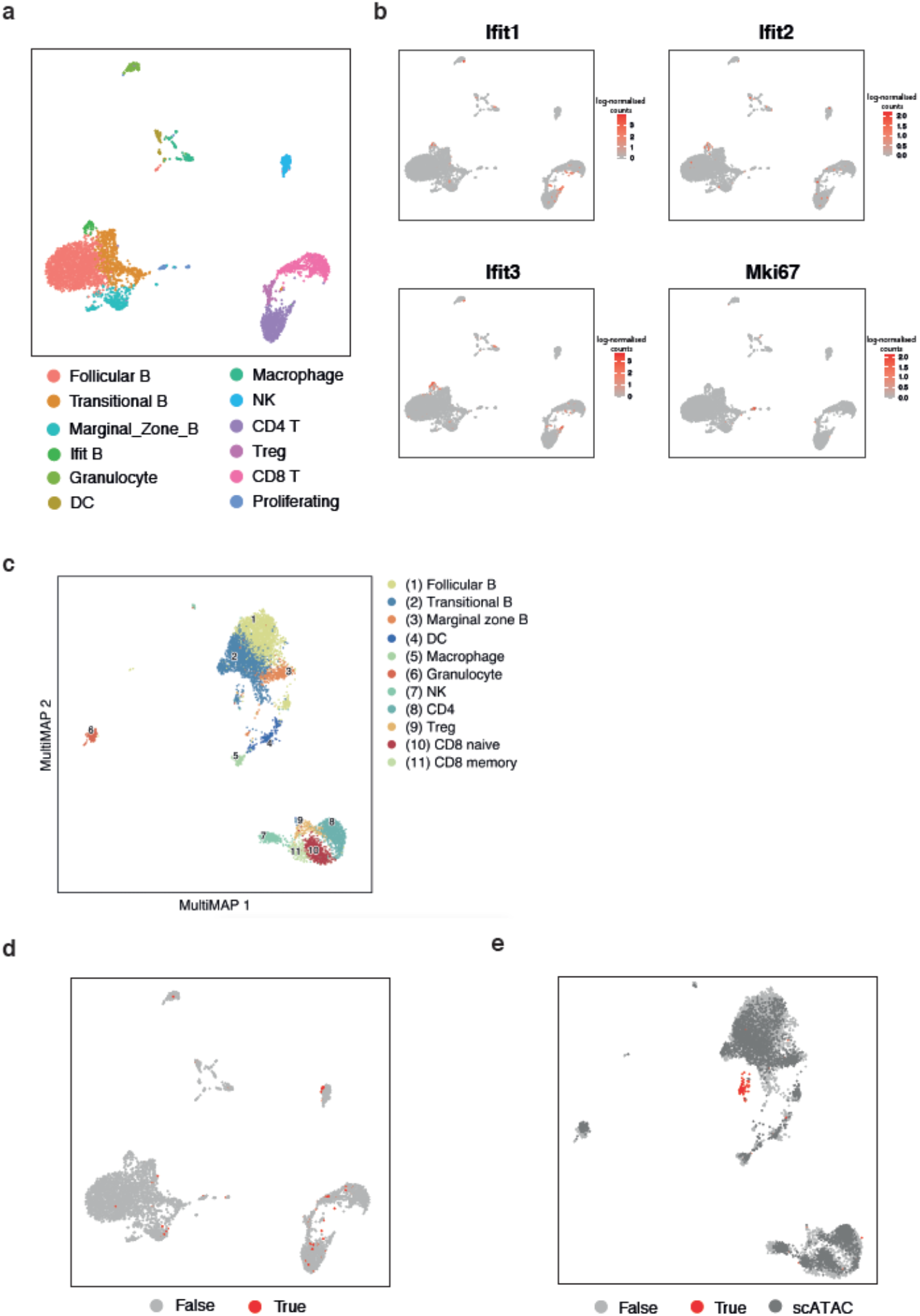
Mouse spleen scRNA-seq and scATAC-seq data. **a**. UMAP visualization of the mouse spleen scRNA-seq data (n=1) colored by the identified cell types. **b**. UMAP visualisation of expression levels of Ifit family genes associated with interferon response, upregulated in one specific B cell subpopulation, and the proliferation marker Mki67. **c**. MultiMAP visualization of the integrated scRNA-seq and scATAC-seq mouse spleen data (n=1) colored by the jointly identified clusters. **d, e**. UMAP (d) and MultiMAP (e) visualizations of the mouse spleen data showing cells identified as doublets (labelled “True”) using an independent pipeline (Scrublet). The MultiMAP visualisation leads to these artifactual data points being clustered in one group, highlighting the power of this method to visualise and separate data.

**Extended Data Figure 4.**
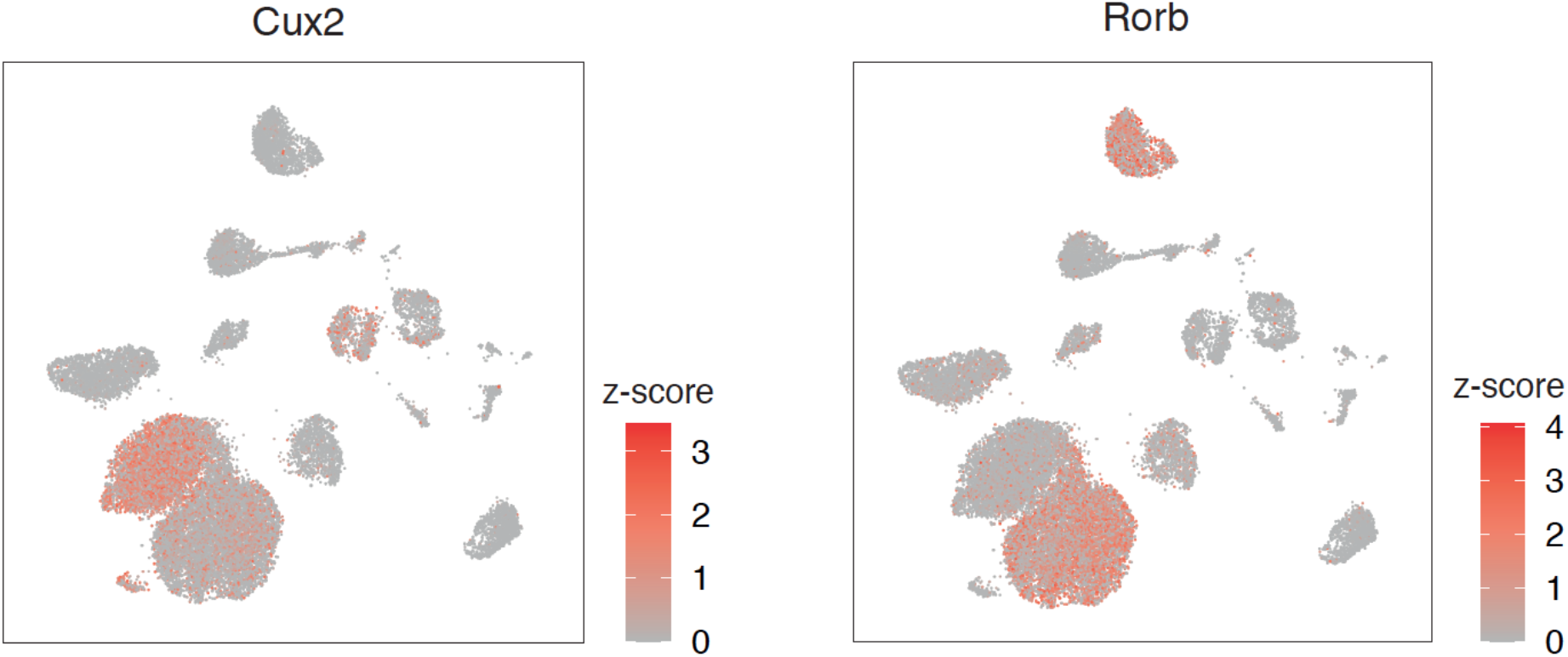
Marker genes of the L4 cluster identified in the scRNA-seq and STARmap integration. MultiMAP visualisation of log-transformed gene expression of markers associated with L4 neurons. The MultiMAP integration identified L4 cells in the scRNA-seq data previously annotated as L5 neurons.

**Extended Data Figure 5.**
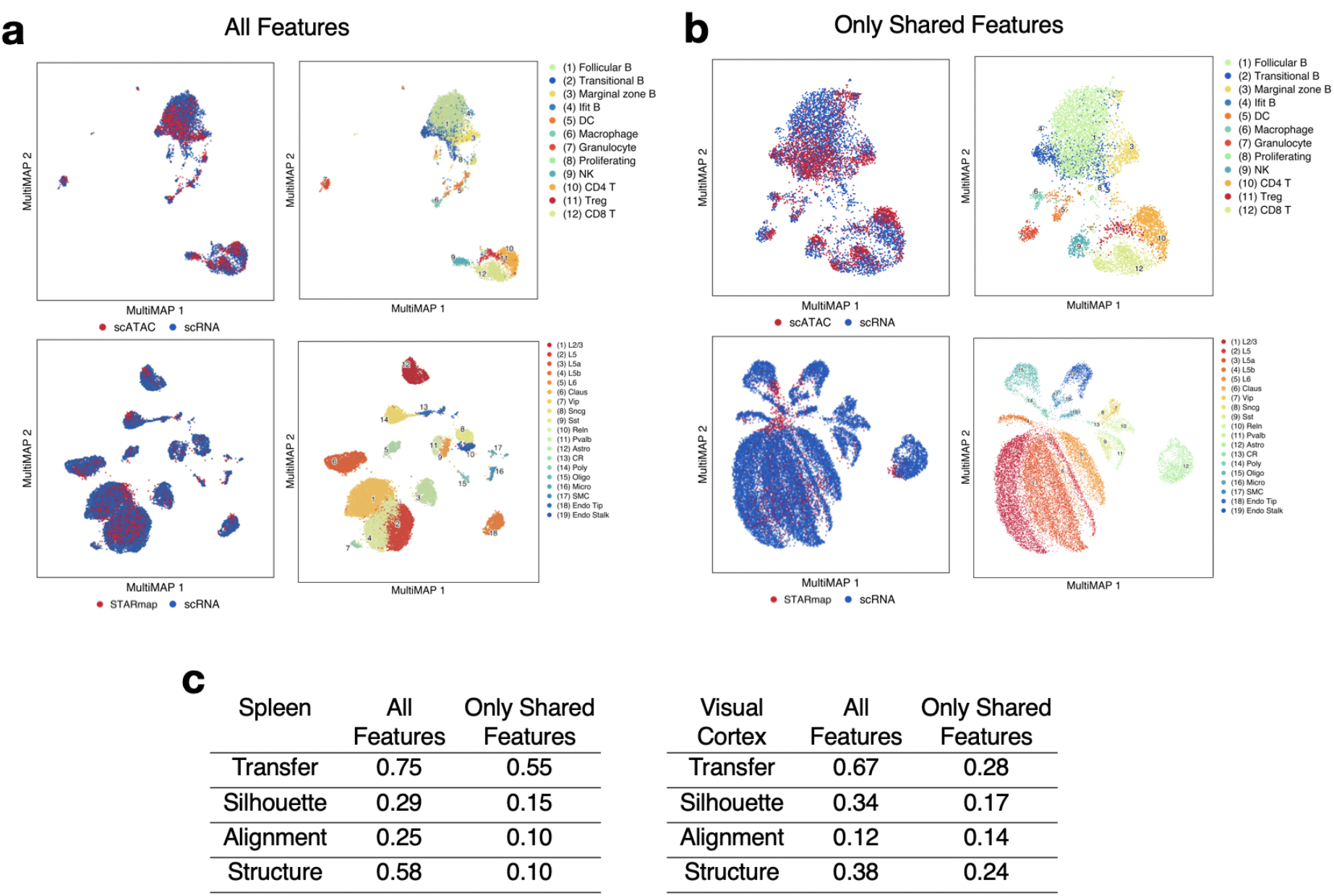
MultiMAP integration with all features vs. only shared features in the spleen scRNA-seq + scATACseq, and visual cortex STARmap + scRNAseq datasets. **a.** MultiMAP embeddings using all genes present in each dataset (intended use of MultiMAP). **b**. MultiMAP embeddings using only genes shared by all datasets in each integration. **c**. Comparison of the MultiMAP integration with all features vs. only shared features --in terms of transfer learning accuracy (“Transfer”), separation of cell type clusters as quantified by Silhouette coefficient (“Silhouette”), mixing of different datasets as measured by fraction of nearest neighbours that belong to a different dataset (“Alignment”), and preservation of high-dimensional structure as measured by the Pearson correlation between distances in the high-and low-dimensional spaces (“Structure”).

**Extended Data Figure 6.**
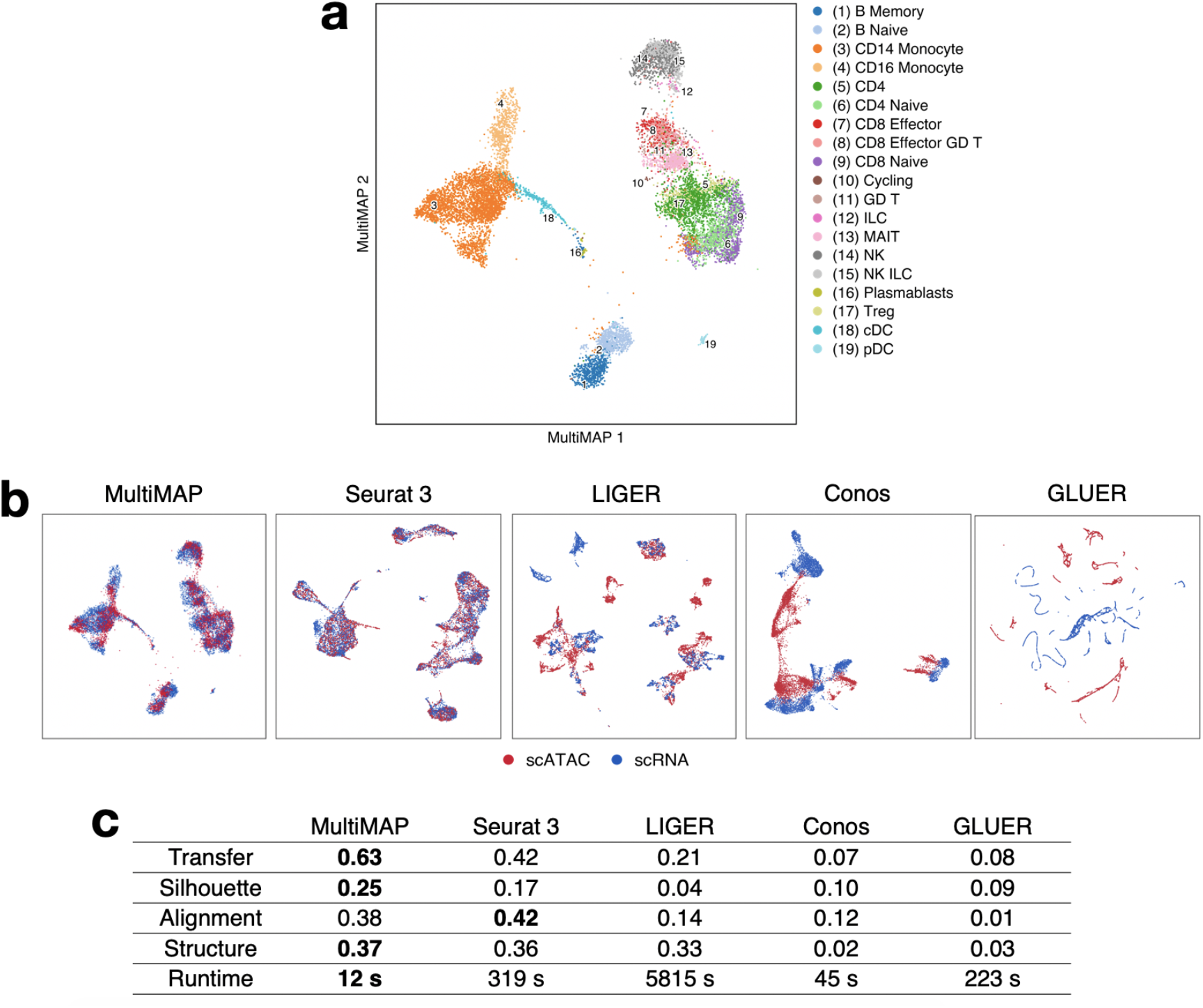
Benchmarking MultiMAP using paired PBMCs. **a**. MultiMAP visualization of the Multiome RNA+ATAC PBMCs, colored by independently annotated cell type. **b**. Embeddings produced by alternative integration strategies, colored by omic technology. **c**. Comparison of each method in terms of transfer learning accuracy (“Transfer”), separation of cell type clusters as quantified by Silhouette coefficient (“Silhouette”), mixing of different datasets as measured by fraction of nearest neighbours that belong to a different dataset (“Alignment”), preservation of high-dimensional structure as measured by the Pearson correlation between distances in the high-and low-dimensional spaces (“Structure”), and runtime.

**Extended Data Figure 7.**
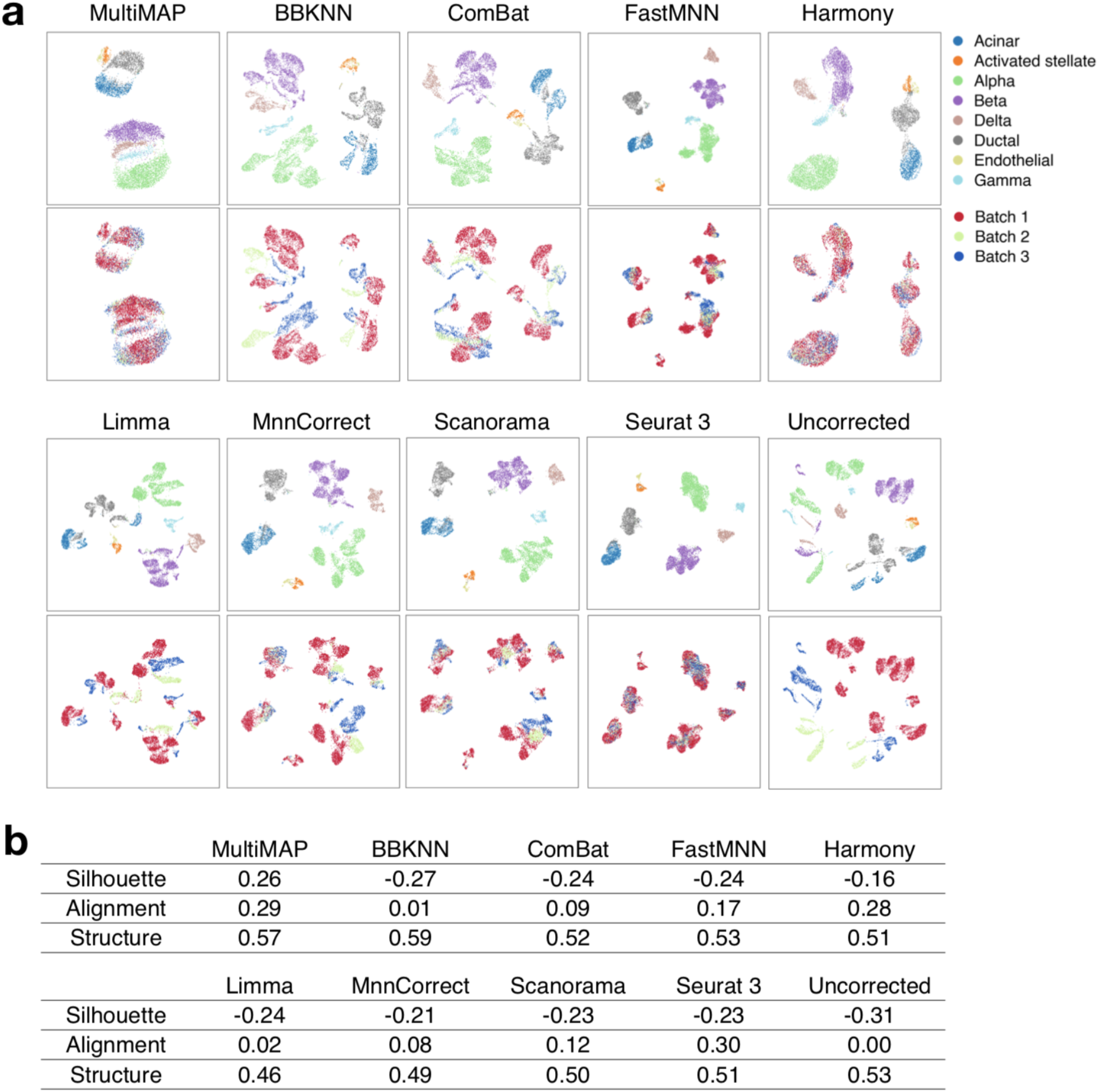
Benchmarking MultiMAP against batch correction methods. **a**. Embeddings returned by MultiMAP and batch correction methods on three scRNA-seq pancreas datasets. **b**. Comparison of separation of cell type clusters as quantified by Silhouette coefficient (“Silhouette”), mixing of different datasets as measured by fraction of nearest neighbours that belong to a different dataset (“Alignment”), preservation of high-dimensional structure as measured by the Pearson correlation between distances in the high-and low-dimensional spaces (“Structure”), and runtime.

**Extended Data Figure 8.**
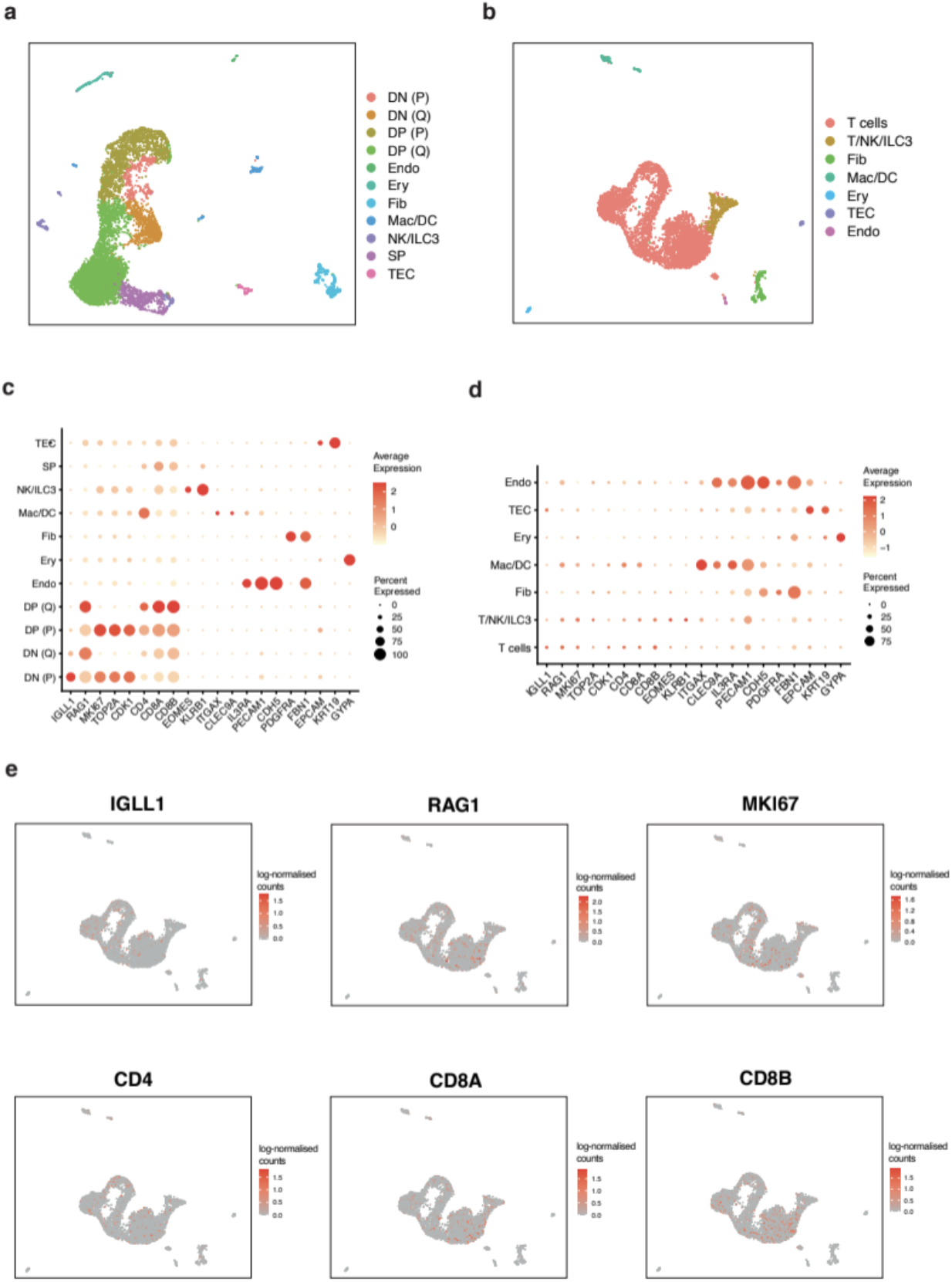
Fetal thymus scRNA-seq and scATAC-seq data. **a**. UMAP visualisation of the fetal thymus scRNA-seq data (n=1) colored by identified cell types shows the same cell types as previously published^31^. **b**. UMAP visualisation of the fetal thymus scATAC-seq data (n=1) colored by the identified cell types. **c**. Dot plot showing the z-score of the mean log-transformed expression level of marker genes. **d**. Dot plot showing the z-score of the mean log-transformed gene activity scores of marker genes, showing not very clear separation of T cells clusters in the scATAC-seq data. **e**. UMAP visualisation of log-transformed gene activity scores of markers for specific T cell subpopulations, showing that the scATAC-seq dataset does not separate well the T cell clusters.

**Extended Data Figure 9.**
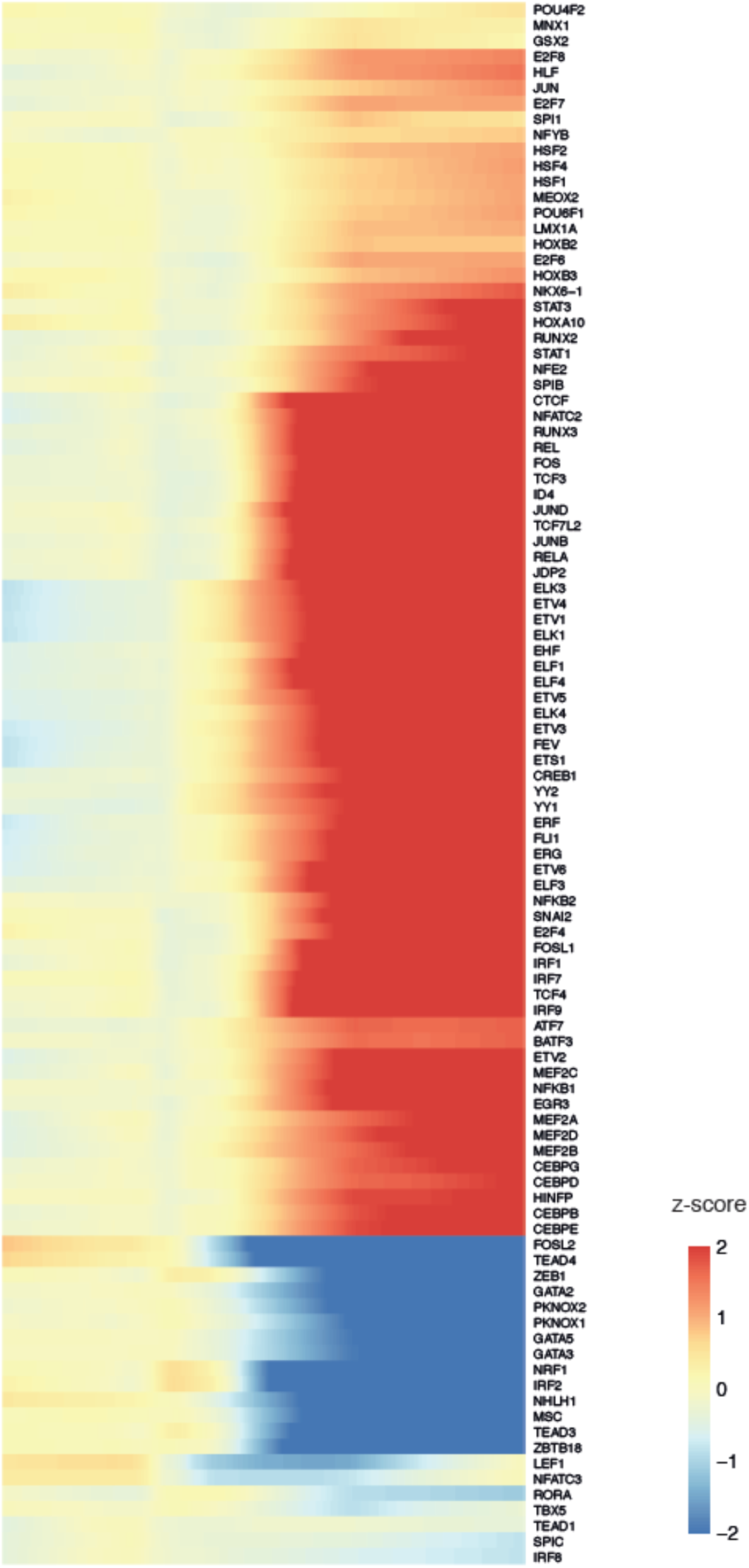
Chromatin accessibility of transcription factor binding sites. Smoothed heatmaps of the z-score of motif accessibility of the top 100 most variable transcription factor binding sites over pseudotime. The TF binding sites that varied most in time show changes in accessibility at the transition between the late DN and early DP stage of differentiation.

## Methods

### MultiMAP

MultiMAP (Figure 1) is a new approach for the integration and dimensionality reduction of multimodal data based on a framework of Riemannian geometry and algebraic topology. MultiMAP takes as input any number of datasets of potentially differing dimensions. The datasets take the form **X**^i^, *i* =1,2,…, with **x** _j_^i^ ∈ R^Di^ being the *j*’th point in dataset **X**^i^. MultiMAP recovers geodesic distances on a single latent manifold *M* on which all of the data is uniformly distributed. The geodesic distances are calculated between data points of the same dataset by normalizing distances in each dataset’s ambient space X^ii^ with respect to a neighborhood distance specific to the dataset, and between data points of different datasets by normalizing distances between the data in a shared ambient space X^ij^ with respect to a neighborhood distance specific to the shared feature space. When integrating multi-omics data with MultiMAP, the ambient spaces are the PC components of each dataset’s full feature space and of the shared feature space(s). These neighborhood distances are the radius of a constant-radius ball *B* on *M*. These distances are then used to construct a neighborhood graph (MultiGraph) on the manifold. Finally the data and manifold space are projected into a low-dimensional embedding space by minimizing the cross entropy of the graph in the embedding space with respect to the graph in the manifold space. Specifically, this optimization minimizes cross entropy of a fuzzy set–representation (*ν*, {**x**_j_ ^i^}) of the graph in the embedding space with respect to a fuzzy set–representation (*μ*, {**x**_j_ ^i^}) of the graph in the manifold space. MultiMAP allows the user to modify the weight *ω*^i^ of each dataset in the cross entropy loss, allowing the user to modulate the contribution of each dataset to the layout. Integrated analysis can be performed on the embedding or the graph, and the embedding also provides an integrated visualization. An extended description of MultiMAP, including mathematical background, is in the Supplementary information.

### Synthetic Data

MultiMAP was applied to two synthetic examples of multimodal data, in order to study the technique in a controlled setting.

The first synthetic setting is schematized in Figure 2a. This setting consists of one dataset (**X**^1^) of 10,000 points sampled randomly from the canonical 3D “Swiss roll” surface (generated with sklearn in Python), and a second dataset (**X**^2^) of 10,000 points sampled randomly from a 2D rectangle. The two datasets can be considered multimodal data because they have different feature spaces but describe a similar rectangular manifold. In addition, we are given the position along the manifold of 1% of the data. Distances between data in the different datasets are calculated for 1% of the data as the absolute differences between these positions. These distances are supplied to MultiMAP. The purpose of this setting is to determine if MultiMAP can integrate data in a nonlinear fashion and operate on datasets of different dimensionality.

The second synthetic setting is schematized in Figure 2c. This setting consists of two datasets based on the MNIST database^34^ which comprises 70,000 28×28 pixel grayscale images of handwritten digits 0-9. The first dataset (**X**^1^) consists of the 28×15 pixel left half of each of images flattened into a 420 dimensional vector. The second dataset (**X**^2^) consists of the 28×15 pixel right half of each of 70,000 digit images, also flattened into a 420 dimensional vector. Added to the first dataset is Gaussian noise with a mean of zero and a standard deviation equal to the maximum pixel value. The two halves overlap by a 28×2 pixel region. Distances between data in the different datasets are calculated in this shared space and supplied to MultiMAP. The two datasets can be considered multimodal because they have different feature spaces but describe a similar population of digit images. The purpose of this setting is to determine if MultiMAP can effectively leverage features unique to certain datasets. The thin overlapping region of the two halves is not enough information to create a good embedding of the data. Many distinct digits are similar in this thin central sliver, and hence they should cluster together in the feature space of the two pixel overlap. Indeed, in a UMAP projection of the data in the shared feature space of this overlap, the clusters of different digits are not as well separated as in the UMAP projections of each half (Figure 2c). A multimodal integration strategy that effectively leverages all features would use the features unique to each half to separate different digits, and the shared space to bring the same digits from each dataset close together.

### Acquisition and processing of human fetal thymic tissue

The tissue sample used for this study was obtained with written informed consent from the participant in accordance with the guidelines in The Declaration of Helsinki 2000. The human fetal tissue was obtained from the MRC/Wellcome Trust-funded Human Developmental Biology Resource (HDBR, http://www.hdbr.org) with appropriate maternal written consent and approval from the Newcastle and North Tyneside NHS Health Authority Joint Ethics Committee (08/H0906/21+5). HDBR is regulated by the UK Human Tissue Authority (HTA; www.hta.gov.uk) and operates in accordance with the relevant HTA Codes of Practice.

The developmental age was estimated from measurements of foot length and heel-to-knee length, and compared against a standard growth chart^35^. A piece of skin was collected from every sample for Quantitative Fluorescence-Polymerase Chain Reaction analysis using markers for the sex chromosomes and the following autosomes: 13, 15, 16, 18, 21, 22. The sample was of normal karyotype.

The tissue was processed immediately after isolation using enzymatic digestion. Tissue was transferred to a sterile 10mm^2^ tissue culture dish and cut into <1mm^3^ segments before being transferred to a 50mL conical tube. Tissues were digested with 1.6mg/mL collagenase type IV (Worthington) in RPMI (Sigma-Aldrich) supplemented with 10%(v/v) heat-inactivated fetal bovine serum (FBS; Gibco), 100U/mL penicillin (Sigma-Aldrich), 0.1mg/mL streptomycin (Sigma-Aldrich), and 2mM L-glutamine (Sigma-Aldrich) for 30 minutes at 37°C with intermittent shaking. Digested tissue was passed through a 100µm filter, and cells collected by centrifugation (500g for 5 minutes at 4°C). Cells were treated with 1X red blood cells (RBC lysis buffer (eBioscience) for 5 minutes at room temperature and washed once with a flow buffer (PBS containing 5%(v/v) FBS\ and 2mM EDTA) prior to cell counting. For scATAC-seq, cells were taken forward for nuclei isolation following 10X Genomics guidelines. Briefly, cells were centrifuged (300g for 5 minutes), added the lysis buffer (Tris-HCl (pH 7.4) 10mM; NaCl 10Mm; MgCl2 3mM; Tween-20 0.1%; NP-40 0.1%; Digitonin 0.01%; BSA 1%) and incubated on ice for 3 minutes (time optimized for thymus). Following the incubation, cells were washed (Tris-HCl (pH 7.4) 10mM; NaCl 10Mm; MgCl2 3mM; BSA 1%; Tween-20 0.1%) and centrifuged (300g for 5 minutes) and nuclei were resuspended in Diluted Nuclei Buffer (10X Genomics). Isolated nuclei were high-quality with well-resolved edges and no evidence of blebbing. The final nuclei concentration was determined prior to loading using a hemocytometer.

### Single-cell RNA and ATAC sequencing of human thymus

scRNA-seq targeting 5,000 cells per sample was performed using the Chromium Controller (10x Genomics). Single-cell cDNA synthesis, amplification, and sequencing libraries were generated using the Single Cell 5’ Reagent Kit following the manufacturer’s instructions. The libraries from up to eight loaded channels were multiplexed together and sequenced on an Illumina HiSeq 4000.

scATAC-seq targeting 5,000 cells was performed using Chromium Single Cell ATAC Library and Gel Bead kit (10x Genomics). The libraries from up to eight loaded channels were multiplexed together and sequenced on an Illumina HiSeq 4000.

### Computational processing and analysis of the human fetal thymus single cell genomics data

scRNA-seq data were aligned and quantified using the Cell Ranger Single-Cell Software Suite (version 2.0, 10x Genomics) against the GRCh38 human reference genome provided by Cell Ranger. The scRNA-seq data was preprocessed using Seurat 3. Cells with fewer than 500 detected genes and more than 10% mitochondrial gene expression content were removed. Ribosomal genes, cell cycle genes^31^ and genes associated with dissociation-induced effects^36^ were removed. Clusters were identified using a community identification algorithm as implemented in the Seurat ‘FindClusters’ function, using 30 principal components (PCs) and annotated using canonical cell-type markers from^31^.

The scATAC-seq data was aligned and preprocessed using CellRanger (10x Genomics). SnapATAC^37^ was used for quality control, preprocessing, and generating cell-by-bin and log-normalized gene activity matrices. The binarized cell-by-bin matrix was used as input for term frequency-inverse document frequency (TF-IDF) weighting, using term frequency and smoothed inverse document frequency as the weighting scheme. Weighted data were reduced to 30 dimensions using singular-value decomposition (SVD). Clustering and UMAP visualization were performed using Seurat 3. chromVar^38^ was used to discover transcription factor dynamics and variation in their motif accessibility.

The 50 dimension reduced scATAC-seq and the 50 dimension reduced scRNA-seq data were supplied as input to MultiMAP. A shared feature space with both the scATAC-seq and scRNA-seq was constructed by removing genes from each dataset that were not present in the other, and then reducing the space to 50 dimensions using PCA. This shared space was supplied as input to MultiMAP, allowing the calculation of distances between cells from different datasets. The parameters of MultiMAP were all set to their default values, including the weight parameter for the scRNA-seq set to 0.8 and for ATAC-seq set to 0.2, on account of the higher-quality scRNA-seq.

The Leiden algorithm^39^ was applied directly to the MultiGraph to jointly cluster all cells. The clusters were then annotated using canonical cell-type markers from ^31^. Diffusion pseudotime (DPT)^40^ was used for trajectory inference. The MultiGraph was supplied as input to the DPT function in SCANPY^41^. DPT was performed only on cells annotated as T cells. Cells were removed if they were positioned away from T cell clusters and close to Fibroblasts and Erythrocytes on the MultiMAP plot, as this likely indicated that they were incorrectly annotated. tradeSeq^42^ was used to identify genes whose expression changes significantly along the trajectory.

### Acquisition and processing of human PBMCs

PBMCs from two donors were acquired from a LeukoLab (Clinical division of AllCells). Frozen PBMC samples were thawed quickly at 37 °C in a water bath. Two pools made for technical duplicates with ∼500,000 cells for each donor per pool (50/50). Nuclei isolation, transposition, ATAC-seq and Gene Expression (GEX) sequencing libraries construction performed according to the manufacturer’s Demonstrated protocol (CG000365 Rev A; 10X Genomics) and Next GEM Single Cell Multiome ATAC and Gene Expression user guide (CG000338 Rev A; 10X Genomics). One lane per pool with a 3,000 targeted nuclei recovery was loaded on a Chromium Next GEM Chip J. ATAC-seq and GEX indexed libraries were sequenced on a NovaSeq 6000 SP Flowcell according to the 10X Genomics recommendations, aiming for a minimum of 50,000 PE reads per cell for both types (ATAC-Seq and GEX) libraries.

### Computational processing and analysis of the human PBMCs Multiome ATAC+RNA data

snRNA-seq and snATAC-seq data were aligned and quantified using the Cell Ranger ARC suite (10x Genomics) against the GRCh38 human reference genome provided by Cell Ranger. The snRNA-seq data was preprocessed using Seurat 3. Cells with fewer than 500 detected genes and more than 20% mitochondrial gene expression content were removed. Clusters were identified using a community identification algorithm as implemented in the Seurat ‘FindClusters’ function, using 30 principal components (PCs) and annotated using canonical cell-type markers.

SnapATAC^37^ was used for quality control, preprocessing, and generating cell-by-bin and log-normalized gene activity matrices for the snATAC-seq data. The binarized cell-by-bin matrix was used as input for term frequency-inverse document frequency (TF-IDF) weighting, using term frequency and smoothed inverse document frequency as the weighting scheme. Weighted data were reduced to 30 dimensions using singular-value decomposition (SVD). Clustering and UMAP visualization were performed using Seurat 3. We used the The 50 dimension reduced snATAC-seq and the 50 dimension reduced snRNA-seq data were supplied as input to MultiMAP. A shared feature space with both the snATAC-seq and snRNA-seq was constructed by removing genes from each dataset that were not present in the other, and then reducing the space to 50 dimensions using PCA. This shared space was supplied as input to MultiMAP, allowing the calculation of distances between cells from different datasets. The parameters of MultiMAP were all set to their default values, including the weight parameter for the snRNA-seq set to 0.8 and for snATAC-seq set to 0.2, on account of the higher-quality snRNA-seq.

### Single-cell RNA sequencing of mouse spleen and data processing

The mice were maintained under specific pathogen-free conditions at the Wellcome Trust Genome Campus Research Support Facility (Cambridge, UK). These animal facilities are approved by and registered with the UK Home Office. All procedures were in accordance with the Animals (Scientific Procedures) Act 1986. The protocols were approved by the Animal Welfare and Ethical Review Body of the Wellcome Trust Genome Campus.

The spleen from a 6-month-old C57BL/6Jax mouse was removed. The splenocytes were isolated by passing the spleen through a 70 µm cell strainer (Fisher Scientific 10788201) into 30 m ice-cold 1X DPBS (Thermo Fisher 14190169) with 2 mM EDTA and 0.5% (w/v) BSA (Sigma A9418) using the plunger of a 2-ml syringe. Cells were spun down at 500 g for 7 minutes at 4 degree. Then the supernatant was removed, and the cell pellet resuspended in 5 ml 1X RBC lysis buffer (Thermo Fisher 00-4300-54). The cell suspension was vigorously vortexed for 5 seconds and left on the bench for 5 minutes to lyse the red blood cells. Then 45 ml ice-cold 1X DPBS was added, and cells were spun down at 500 g for 7 minutes at 4 degrees. The supernatant was removed, and 30 ml ice-cold 1X DPBS with 0.1% BSA was used to resuspend the cell pellet. The cell suspension was passed through a Miltenyi 30 μm Pre-Separation Filter (Miltenyi 130-041-407), and the cell number was determined using the C-chip counting chamber (VWR DHC-N01). The cells were spun down again, and the cell pellet resuspended in ice-cold 1X DPBS with 0.1% BSA to reach a concentration of 1,000,000 cells per ml. The splenocytes were then loaded on the 10x Chromium Controller, aiming to recover ∼ 5000 cells (Targeted Cell Recovery 5000 cells). cDNA and a sequencing library were made according to 10x Single Cell 3’ Reagent Kits v2 manual. The library was sequenced on an Illumina HiSeq 4000 machine.

The resulting scRNA-seq data were preprocessed using CellRanger (10x Genomics) and downstream analysis were performed using the Seurat 3 workflow. Cells with fewer than 200 detected genes and more than 10% mitochondrial gene expression content were filtered out. Downstream analyses such as normalization, clustering and visualization were performed using Seurat 3. Clusters were identified using the community identification algorithm as implemented in the Seurat ‘FindClusters’ function, using 20 PCs. Clusters were annotated using canonical cell-type markers from the original study^15^. Scrublet^43^ was used for doublet detection.

### Acquisition and processing of previously published datasets

The mouse spleen scATAC-seq data was obtained from ArrayExpress (E-MTAB-6714) and preprocessed using the code provided by Chen et al. ^15^(https://github.com/dbrg77/plate_scATAC-seq). Briefly, reads from all cells were merged, and open chromatin regions were identified by peak calling with MACS2 ^44^. Latent semantic indexing analysis was used for dimensionality reduction of the resulting cell-by-bin matrix. The binary cell-by-bin accessibility was used as input for TF-IDF weighting, using term frequency and smoothed inverse document frequency as the weighting scheme. Weighted data were reduced to 50 dimensions using SVD. SnapATAC^37^ was used to generate gene activity count matrices, which were then log-normalized. The 50 dimension reduced accessibility of the scATAC-seq and the 50 dimension reduced gene expression of the scRNA-seq data were supplied as input to MultiMAP. The 50-dimension reduced accessibility of the scATAC-seq and the 50-dimension reduced gene expression of the scRNA-seq data were supplied as input to MultiMAP. A shared feature space with both the scATAC-seq and scRNA-seq was constructed by removing genes from each dataset that were not present in the other, and then reducing the space to 50 dimensions using PCA. This shared space was supplied as input to MultiMAP, allowing the calculation of distances between cells from different datasets. The parameters of MultiMAP were all set to their default values, including the weight parameter for the scRNA-seq set to 0.8 and for ATAC-seq set to 0.2 due to the higher quality scRNA-seq. The Leiden algorithm was applied directly to the MultiGraph to jointly cluster all cells. Harmonic function-based node classification was performed directly on the MultiGraph to predict cell types of the scATAC-seq cells given the cell types of the scRNA-seq cells^45^.

Human hematopoiesis scRNA-seq and scATAC-seq data were downloaded from https://github.com/GreenleafLab/MPAL-Single-Cell-2019. The scRNA-seq consists of 6 experimental batches, and the scATAC-seq consists of 10 experimental batches. Severe batch effects were observed, so this data was considered to consist of 16 separate datasets for the integration with MultiMAP. The scRNA-seq data was preprocessed using Seurat 3, and each batch was log-normalized and reduced to 50 dimensions with PCA. The cell-by-bin peak accessibility was used as provided by the authors. The binary cell-by-bin accessibility was used as input for TF-IDF weighting, using term frequency and smoothed inverse document frequency as the weighting scheme. Separately for each batch, the weighted data were reduced to 50 dimensions using SVD. Gene activities of the ATAC data were calculated using Cicero^46^ and log-normalized. To integrate all of the data at once, all 16 datasets were provided as input to MultiMAP in the form of the 50 dimension reduced accessibility data of the scATAC-seq and the 50 dimension reduced gene expression of the scRNA-seq. Shared feature spaces containing two datasets were constructed by removing genes from each of the datasets that were not present in the other, and then reducing the space to 50 dimensions using PCA. These shared spaces were supplied as input to MultiMAP to calculate distances between cells from different datasets. The parameters of MultiMAP were all set to their default values, including the weight parameter for the scRNA-seq set to 0.8 and for ATAC-seq set to 0.2 due to the higher-quality scRNA-seq data.

scRNA-seq data of the mouse frontal cortex acquired with Drop-seq was obtained from dropviz.org. STARmap data of the mouse visual cortex was downloaded from https://www.starmapresources.com/data/. Each dataset was separately preprocessed with Seurat 3^11^, log-normalized, and reduced to 50 dimensions with PCA. Both 50 dimensional reduced datasets were supplied as input to MultiMAP. A shared feature space with both the STARmap and scRNA-seq data was constructed by removing genes from each dataset that were not present in the other, and then reducing the space to 50 dimensions using PCA. This shared space was supplied as input to MultiMAP to calculate distances between cells from different datasets. The parameters of MultiMAP were all set to their default values, including the weight parameter for the scRNA-seq set to 0.8 and for Drop-seq set to 0.2, on account of higher-quality, tighter clusters generally observed in the scRNA-seq.

scRNA-seq, scATAC-seq, and snmC-seq data from the mouse primary cortex^20^ was downloaded from the Neuroscience Multi-omics Archive (NeMO). The scRNA-seq was preprocessed using Seurat 3, log-normalised, and reduced to 50 dimensions with PCA. The binary cell-by-bin accessibility and gene activity count matrix of the scATAC-seq were obtained with SnapATAC^37^. The gene activity count data was log-normalized. Latent semantic indexing analysis was used for dimensionality reduction of the scATAC-seq accessibility. The binary cell-by-bin accessibility was used as input for TF-IDF weighting, using term frequency and smoothed inverse document frequency as weighting scheme. Weighted data were reduced to 50 dimensions using SVD. The DNA methylation data was preprocessed as described in ^47^, using the provided scripts. Briefly, after mapping, the methyl-cytosine counts and total cytosine counts were calculated in two sets of genome regions for each cell: the non-overlapping 100 kb bins tiling the mm10 genome, which was used for dimensionality reduction, and gene body regions ± 2 kb, which is used for the joint alignment. Posterior mCH and mCG rates were calculated based on beta-binomial distribution for the non-overlapping 100kb bins matrix. The top 3000 highly variable features were taken and the data was reduced to 50 dimensions with PCA. Because gene body mCH proportions are negatively correlated with gene expression level, the direction of the methylation data was reversed by subtracting all values from the maximum methylation value^12^. The 50 dimensional reduced scRNA-seq, scATAC-seq, and snmC-seq were supplied as input to MultiMAP. Shared feature spaces containing each pair of two datasets and all three datasets together were constructed by removing genes from each of the datasets that were not present in the other, and then reducing the space to 50 dimensions using PCA. These shared spaces were supplied as input to MultiMAP, allowing the calculation of distances between cells from different datasets. The parameters of MultiMAP were all set to their default values. The weight parameter for the scRNA-seq set to 0.8 and for the other omics set to 0.2, on account of the higher-quality scRNA-seq data.

### Benchmarking

Benchmarking of MultiMAP, Seurat 3, LIGER, Conos and GLUER was performed using a variety of multi-omic data including the scRNA-seq and scATAC-seq data of the spleen, scRNA-seq and STARmap of the visual cortex, and the scRNA-seq, scATAC-seq, and snmC-seq of the primary cortex. These datasets were chosen because they all have cell type annotations supplied in their original publications, which was used to independently validate the integration.

The scRNA-seq and STARmap data was log-normalised using Seurat 3 and then used as an input for all integration methods, except GLUER where the raw data was used as an input and preprocessed using the SCANPY workflow. The scATAC-seq data was preprocessed as described above and the log-normalised gene activity matrix was used as an input for all integration methods. Seurat 3, LIGER, Conos and GLUER were executed as detailed in their tutorials, with all parameters set to their default values. Latent Semantic Indexing was used as the dimensionality reduction technique for the scATAC-seq data for weighting anchors in Seurat 3. CCA was used as the dimensionality reduction technique for the scRNA-seq and STARmap data for weighting anchors in Seurat 3.

A diversity of performance metrics was used. After integration, label transfer of the cell type annotations from the scRNA-seq to each other omic was performed by setting the cell type of a query cell to the most frequent type among its 5 nearest labeled neighbors. The balanced accuracy of the label transfer (“Transfer”) was calculated using the annotations from the original publications as the ground truth. A high accuracy indicates that the same cell types from different modalities are near each other in the integrated embedding. After integration, the average Silhouette score^48^ (“Silhouette”‘) across all cells was calculated using the cell type annotations from the original publications as the cluster labels. We note that the Silhouette score is not affected by the number of clusters as we use the same cell type labels, and hence number of clusters, for each integration method. A higher Silhouette score indicates the embedding is better separating distinct cell types. The degree of alignment (“Alignment”‘) of the different datasets in the integrated embedding was calculated as the proportion of each cell’s 5 nearest neighbors that originated in a different dataset, averaged over all cells. This metric was also used in ^12^. A higher value of the alignment score indicates that the different datasets are more evenly mixed in the integrated embedding. The degree to which the embedding preserves the high-dimensional structure (“Structure”) of each dataset was calculated as the Pearson correlation between all pairwise distances in the high-dimensional spaces and the corresponding distances in the embedding. A higher correlation indicates that the embedding is more faithful to the high-dimensional structure. All of these performance metrics were also calculated in the shared feature space of the datasets to be integrated, to get baseline values of the metrics prior to the application of any integration strategy.

The wall-clock runtime of each method on each dataset was recorded. Additionally, to characterize the runtimes of the methods on a wide range of dataset sizes, the integration methods were run on datasets ranging from 1,000 to 500,000 cells. To produce these datasets we subsampled the mouse primary cortex scRNA-seq and scATAC-seq data^20^ using geometric sketching^33^. The datasets were subsampled so that there are equal number of cells in the scRNA-seq and scATAC-seq data until 100,000 cells. Since the scATAC-seq data had 81,196 cells in total, for the 500,000 cells comparison, we used an scRNA-seq of 418,804 cells. All methods were run with 3.1 GHz Intel i7 cores and 218 GB RAM.

## References

1. Becht, E. et al. Dimensionality reduction for visualizing single-cell data using UMAP. Nat. Biotechnol. (2018) doi:10.1038/nbt.4314.

2. Stoeckius, M. et al. Simultaneous epitope and transcriptome measurement in single cells. Nature Methods vol. 14 865–868 (2017).

3. Peterson, V. M. et al. Multiplexed quantification of proteins and transcripts in single cells. Nature Biotechnology vol. 35 936–939 (2017).

4. Klemm, S. L., Shipony, Z. & Greenleaf, W. J. Chromatin accessibility and the regulatory epigenome. Nat. Rev. Genet. 20, 207–220 (2019).

5. Karemaker, I. D. & Vermeulen, M. Single-Cell DNA Methylation Profiling: Technologies and Biological Applications. Trends Biotechnol. 36, 952–965 (2018).

6. Mayr, U., Serra, D. & Liberali, P. Exploring single cells in space and time during tissue development, homeostasis and regeneration. Development 146, (2019).

7. Regev, A. et al. The Human Cell Atlas. Elife 6, (2017).

8. HuBMAP Consortium. The human body at cellular resolution: the NIH Human Biomolecular Atlas Program. Nature 574, 187–192 (2019).

9. Efremova, M. & Teichmann, S. A. Computational methods for single-cell omics across modalities. Nat. Methods 17, 14–17 (2020).

10. Lähnemann, D. et al. Eleven grand challenges in single-cell data science. Genome Biol. 21, 31 (2020).

11. Stuart, T. et al. Comprehensive Integration of Single-Cell Data. Cell vol. 177 1888– 1902.e21 (2019).

12. Welch, J. D. et al. Single-Cell Multi-omic Integration Compares and Contrasts Features of Brain Cell Identity. Cell vol. 177 1873–1887.e17 (2019).

13. Lopez, R. et al. A joint model of unpaired data from scRNA-seq and spatial transcriptomics for imputing missing gene expression measurements. arXiv [cs.LG] (2019).

14. GradientBased Learning Applied to Document Recognition. Intelligent Signal Processing (2009) doi:10.1109/9780470544976.ch9.

15. Chen, X., Miragaia, R. J., Natarajan, K. N. & Teichmann, S. A. A rapid and robust method for single cell chromatin accessibility profiling. Nat. Commun. 9, 5345 (2018).

16. Granja, J. M. et al. Single-cell multiomic analysis identifies regulatory programs in mixed-phenotype acute leukemia. Nat. Biotechnol. 37, 1458–1465 (2019).

17. Saunders, A. et al. Molecular Diversity and Specializations among the Cells of the Adult Mouse Brain. Cell 174, 1015–1030.e16 (2018).

18. Wang, X. et al. Three-dimensional intact-tissue sequencing of single-cell transcriptional states. Science 361, (2018).

19. Brodmann, K. Brodmann’s: Localisation in the Cerebral Cortex. (Springer Science & Business Media, 2007).

20. Yao, Z. et al. An integrated transcriptomic and epigenomic atlas of mouse primary motor cortex cell types. 2020.02.29.970558 (2020) doi:10.1101/2020.02.29.970558.

21. Yamawaki, N., Borges, K., Suter, B. A., Harris, K. D. & Shepherd, G. M. G. A genuine layer 4 in motor cortex with prototypical synaptic circuit connectivity. Elife 3, e05422 (2014).

22. Barkas, N. et al. Joint analysis of heterogeneous single-cell RNA-seq dataset collections. Nat. Methods 16, 695–698 (2019).

23. Peng, T., Chen, G. M. & Tan, K. GLUER: integrative analysis of single-cell omics and imaging data by deep neural network. doi:10.1101/2021.01.25.427845.

24. Muraro, M. J. et al. A Single-Cell Transcriptome Atlas of the Human Pancreas. Cell Syst 3, 385–394.e3 (2016).

25. Segerstolpe, Å. et al. Single-Cell Transcriptome Profiling of Human Pancreatic Islets in Health and Type 2 Diabetes. Cell Metab. 24, 593–607 (2016).

26. Baron, M. et al. A Single-Cell Transcriptomic Map of the Human and Mouse Pancreas Reveals Inter-and Intra-cell Population Structure. Cell Syst 3, 346–360.e4 (2016).

27. Chazarra-Gil, R., van Dongen, S., Kiselev, V. Y. & Hemberg, M. Flexible comparison of batch correction methods for single-cell RNA-seq using BatchBench. Nucleic Acids Res. (2021) doi:10.1093/nar/gkab004.

28. Roels, J. et al. Distinct and temporary-restricted epigenetic mechanisms regulate human αβ and γδ T cell development. Nat. Immunol. 21, 1280–1292 (2020).

29. Jia, G. et al. Single cell RNA-seq and ATAC-seq analysis of cardiac progenitor cell transition states and lineage settlement. Nat. Commun. 9, 4877 (2018).

30. Chen, H. et al. Single-cell trajectories reconstruction, exploration and mapping of omics data with STREAM. Nat. Commun. 10, 1903 (2019).

31. Park, J.-E. et al. A cell atlas of human thymic development defines T cell repertoire formation. Science 367, (2020).

32. Hosokawa, H. & Rothenberg, E. V. How transcription factors drive choice of the T cell fate. Nature Reviews Immunology (2020) doi:10.1038/s41577-020-00426-6.

33. Hie, B., Cho, H., DeMeo, B., Bryson, B. & Berger, B. Geometric Sketching Compactly Summarizes the Single-Cell Transcriptomic Landscape. Cell Syst 8, 483–493.e7 (2019).

34. Lecun, Y., Bottou, L., Bengio, Y. & Haffner, P. Gradient-based learning applied to document recognition. Proceedings of the IEEE vol. 86 2278–2324 (1998).

35. Hern, W. M. Correlation of fetal age and measurements between 10 and 26 weeks of gestation. Obstet. Gynecol. 63, 26–32 (1984).

36. van den Brink, S. C. et al. Single-cell sequencing reveals dissociation-induced gene expression in tissue subpopulations. Nat. Methods 14, 935 (2017).

37. Fang, R. et al. Fast and Accurate Clustering of Single Cell Epigenomes Reveals Cis-Regulatory Elements in Rare Cell Types. doi:10.1101/615179.

38. Schep, A. N., Wu, B., Buenrostro, J. D. & Greenleaf, W. J. chromVAR: inferring transcription-factor-associated accessibility from single-cell epigenomic data. Nat. Methods 14, 975–978 (2017).

39. Blondel, V. D., Guillaume, J.-L., Lambiotte, R. & Lefebvre, E. Fast unfolding of communities in large networks. Journal of Statistical Mechanics: Theory and Experiment vol. 2008 P10008 (2008).

40. Haghverdi, L., Büttner, M., Wolf, F. A., Buettner, F. & Theis, F. J. Diffusion pseudotime robustly reconstructs lineage branching. Nat. Methods 13, 845 (2016).

41. Wolf, F. A., Angerer, P. & Theis, F. J. SCANPY: large-scale single-cell gene expression data analysis. Genome Biol. 19, 15 (2018).

42. Van den Berge, K. et al. Trajectory-based differential expression analysis for single-cell sequencing data. Nat. Commun. 11, 1201 (2020).

43. Wolock, S. L., Lopez, R. & Klein, A. M. Scrublet: Computational Identification of Cell Doublets in Single-Cell Transcriptomic Data. Cell Syst 8, 281–291.e9 (2019).

44. Grytten, I. et al. Graph Peak Caller: Calling ChIP-seq peaks on graph-based reference genomes. PLoS Comput. Biol. 15, e1006731 (2019).

45. Zhu, X., Ghahramani, Z. & Lafferty, J. D. Semi-supervised learning using gaussian fields and harmonic functions. in Proceedings of the 20th International conference on Machine learning (ICML-03) 912–919 (2003).

46. Pliner, H. A. et al. Cicero Predicts cis-Regulatory DNA Interactions from Single-Cell Chromatin Accessibility Data. Mol. Cell 71, 858–871.e8 (2018).

47. Kozareva, V. et al. A transcriptomic atlas of the mouse cerebellum reveals regional specializations and novel cell types. doi:10.1101/2020.03.04.976407.

48. Rousseeuw, P. J. Silhouettes: A graphical aid to the interpretation and validation of cluster analysis. Journal of Computational and Applied Mathematics vol. 20 53–65 (1987).

